# Retrospective identification of intrinsic factors that mark pluripotency potential in rare somatic cells

**DOI:** 10.1101/2023.02.10.527870

**Authors:** Naveen Jain, Yogesh Goyal, Margaret C. Dunagin, Christopher J. Cote, Ian A. Mellis, Benjamin Emert, Connie L. Jiang, Ian P. Dardani, Sam Reffsin, Arjun Raj

## Abstract

Pluripotency can be induced in somatic cells by the expression of the four “Yamanaka” factors OCT4, KLF4, SOX2, and MYC. However, even in homogeneous conditions, usually only a rare subset of cells admit reprogramming, and the molecular characteristics of this subset remain unknown. Here, we apply retrospective clone tracing to identify and characterize the individual human fibroblast cells that are primed for reprogramming. These fibroblasts showed markers of increased cell cycle speed and decreased fibroblast activation. Knockdown of a fibroblast activation factor identified by our analysis led to increased reprogramming efficiency, identifying it as a barrier to reprogramming. Changing the frequency of reprogramming by inhibiting the activity of LSD1 led to an enlarging of the pool of cells that were primed for reprogramming. Our results show that even homogeneous cell populations can exhibit heritable molecular variability that can dictate whether individual rare cells will reprogram or not.

## Introduction

The demonstration that pluripotent stem cells can be induced from differentiated somatic cells via ectopic expression of the four “Yamanaka’’ factors (OCT4, KLF4, SOX2, and MYC; OKSM) was a watershed discovery that holds promise for disease modeling and regenerative medicine (Takahashi & Yamanaka, 2006; Yamanaka, 2009). Yet, induction of pluripotency is a highly inefficient process, with only a small percentage of originating cells, often under 0.1%, properly undergoing reprogramming into iPSCs. Often, this low efficiency is ascribed to technical variability in the delivery and stoichiometry of the OKSM factors, but even when the OKSM factors are expressed uniformly from a single promoter in clonally derived cells, these efficiencies remain low, maxing out at around 1-5% for mouse embryonic fibroblasts (Hockemeyer et al., 2008; Maherali et al., 2008; Plath & Lowry, 2011; Polo et al., 2010). One possibility is that intrinsic differences between individual cells *before* the addition of the OKSM factors lead them to either reprogram or not. Profiling the highly-reprogrammable cells could reveal the barriers to reprogramming that are active in the rest of the cells that comprise the majority of the population; however, the identification of these cells and the specific factors within them has remained challenging.

Indeed, even the question of whether the cells that are able to reprogram have a fixed identity remains heavily debated. One set of results argues that there is no intrinsically defined “primed” subpopulation *per se*, and that all cells are equally capable of undergoing reprogramming (Buganim et al., 2012; J. Hanna et al., 2009; J. H. Hanna et al., 2010; Hochedlinger & Jaenisch, 2015). The primary evidence for this model is the observation that all single-cell clones derived from the parental population have a reprogrammable subpopulation (J. Hanna et al., 2009). In such a view, only pseudo-random effects (i.e., unknown environmental factors) can dictate which cells reprogram. Other experiments, however, have provided evidence for the existence of intrinsic differences dictating reprogramming outcomes. Most directly, cells recently derived from a shared single progenitor (i.e., twins) share reprogramming outcomes even when divorced from their original context and separated onto different plates (Pour et al., 2015; Shakiba et al., 2019; Yunusova et al., 2017). These results collectively suggest that reprogramming potential is at least partially encoded by pre-existing differences before OKSM induction in subsets of cells primed to become iPSCs, and these differences are stable enough to exist across cell division in twins.

What, then, are the factors that are associated with this primed state *before* the induction of OKSM, and how do they differ from those that operate *after* induction? Much work over the years has focused on the latter, elucidating the molecular sequence of events between the induction of OKSM in somatic cells and becoming fully reprogrammed as an iPSC. This set of events has been elucidated in great detail via comprehensive analyses of cells undergoing reprogramming. For instance, after induction, cells that reprogram have been shown to exhibit: accelerated cell cycle progression (Babos et al., 2019; S. Guo et al., 2014; J. Hanna et al., 2009; Hu et al., 2019; Smith et al., 2010), an ability to accommodate high rates of transcription (Babos et al., 2019), expression of factors facilitating or marking successful mesenchymal-to-epithelial transition (MET) (Di Stefano et al., 2016; L. Guo et al., 2019; R. Li et al., 2010; X. Liu et al., 2013; Samavarchi-Tehrani et al., 2010; Schwarz et al., 2018), enhanced chromatin accessibility at pluripotency gene loci (Becker et al., 2017; Hussein et al., 2014; Schwarz et al., 2018; Zviran et al., 2018), and expression of factors facilitating the action of the OKSM factors or establishing pluripotency directly (Chronis et al., 2017; Polo et al., 2012; Schwarz et al., 2018; Zviran et al., 2018). However, there is no guarantee that factors responsible for driving the path to pluripotency are the same as those that mark cells primed for reprogramming before induction. A few studies have attempted to show that some features that appear after induction may be present beforehand, such as fast cycling (L. Guo et al., 2019; Utikal et al., 2009) and expression of markers found in developmental progenitors (Nemajerova et al., 2012; Shakiba et al., 2019), even stem cells (Wakao et al., 2011). However, the lack of means to directly and precisely identify primed cells in an unbiased way has limited our knowledge of the factors most critical for priming.

The primary hurdle in identifying these factors is the technical challenge of retrospectively identifying and profiling the initial state of cells based on whether or not they ultimately reprogram into an iPSC. This challenge is compounded by the fact that cells bound to become iPSCs are very rare within the population. Several recently developed clonal barcoding and retrospective characterization methods have now made it possible to connect initial cell state to phenotypic fate with high resolution (Biddy et al., 2018; Emert et al., 2021; Goyal et al., 2021; Umkehrer et al., 2021; Weinreb et al., 2020).

Here, we make use of one such method called Rewind (Emert et al., 2021) that uses a DNA/RNA barcoding strategy to pick out “needle-in-a-haystack” fibroblasts primed to become iPSCs from thousands of nonprimed fibroblasts in a clonally derived population. Using Rewind, we demonstrate the existence of a subset of clonally-derived human fibroblasts primed to become iPSCs upon exposure to OKSM. These primed fibroblasts exhibit an elevated rate of cell cycle progression and have low levels of factors associated with fibroblast activation even before the induction of OKSM. Our results suggest that intrinsic cellular variability can define a reprogrammable state that can persist for multiple cell divisions.

## Results

### Rewind enables retrospective identification and characterization of cells primed for reprogramming into iPSCs

Our goal was to identify markers for cells that are primed for reprogramming. The central challenge was the retrospective identification of the cells that undergo reprogramming. We used a clonal barcoding method called Rewind to explicitly connect pre-existing differences in somatic cells (defining the primed state) with iPSC reprogramming outcomes (Emert et al., 2021). Rewind uses a lentiviral library of barcodes incorporated into the 3’ UTR of GFP, enabling barcodes to stably exist as DNA and mRNA for detection by both single-cell RNA-sequencing and single-molecule RNA FISH imaging. It is particularly well-suited for identifying very rare cells in the population, as is the case with cells primed for reprogramming. In Rewind, after performing barcoding, cells undergo a few divisions resulting in what we refer to as twins, or cells in the same clone with a recent common ancestor. In the case of reprogramming, after separating twins, we immediately transcriptionally profile one split of twins to capture the initial state (i.e., a molecular “carbon copy”), and we reprogram the other split of twins into iPSCs via induction of OKSM. We then sequence the clone barcodes in the resultant iPSCs, use them to identify primed twins in the “carbon copy”, and use their transcriptome profiles to determine which genes’ expression patterns are associated with reprogramming success.

To minimize the potential impact of confounding variables that also may vary between cells, such as the degree of induction of the OKSM factors, we used a clonally-derived, secondary human fibroblast-like cell line (hiF-T) with doxycycline-inducible expression of the OKSM factors (Cacchiarelli et al., 2015) (Figure 1A). The clonal derivation of the line minimized the contribution of genetic differences to reprogramming, and the inducible expression of human telomerase ensured more consistent reprogramming efficiency even after months in culture; these cells displayed less variability in proliferation, senescence and reprogramming efficiency than primary fibroblasts. Furthermore, the OKSM factors are combined in a single cassette, facilitating consistent dosage of OKSM across cells. We observed rapid and relatively homogeneous OKSM induction with low variability—over 99% of cells expressed high levels of OKSM after 48 hours in doxycycline, as determined by measuring *SOX2* and *KLF4* mRNA levels by RNA fluorescence *in situ* hybridization (FISH) with and without doxycycline (Figures S1A-B). Despite the minimization of variability in induction, reprogramming still only occurred in a small fraction of cells (0.01-0.1%).

**Figure 1:**
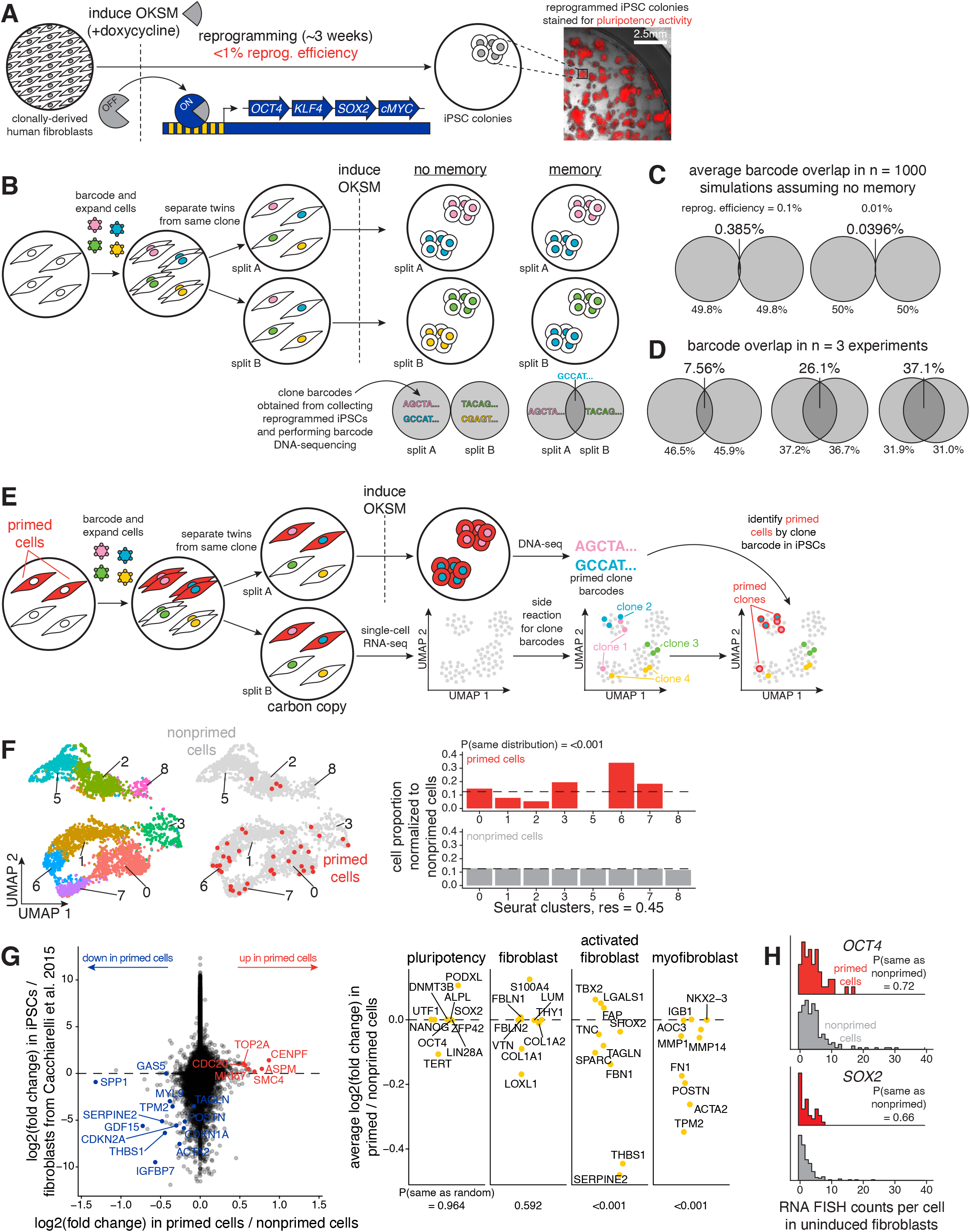
Rewind enables retrospective identification of and gene expression profiling of cells primed for reprogramming into iPSCs before OKSM induction. A. Schematic of reprogramming human inducible fibroblast-like (hiF-T) cells into induced pluripotent stem cells (iPSCs). Addition of doxycycline induces expression of a polycistronic cassette driving stable and stoichiometric expression of the Yamanka factors (OCT4, KLF4, SOX2, and MYC). We stained for alkaline phosphatase activity to identify pluripotent iPSCs by imaging after the reprogramming period of 3-4 weeks. B. Schematic of Rewind for following fates of hiF-T cells. Here, we transduced hiF-T cells at an MOI of ∼1 with our barcode library. After 3-5 cell divisions, we divided the culture into splits (A and B), induced OKSM in each separately, and performed barcode DNA-sequencing in the resulting iPSC colonies. If extrinsic factors alone dictated reprogramming outcomes or if priming did not have memory across cell divisions (i.e., “no memory”), we would expect essentially no overlap in clones forming iPSCs in each split. If intrinsic factors dictated some degree of reprogramming outcomes and if priming had memory across cell divisions (i.e., “memory”), we would expect some degree of overlap in clones forming iPSCs in each split. C. We performed computer simulations to determine the degree of overlap in clones forming iPSCs in each split expected if extrinsic factors alone dictated reprogramming outcomes (i.e., “no memory”). The degree of overlap was simulated for 1000 replicates for reprogramming frequencies of 0.01% and 0.1%, where reprogramming frequency = (number of alkaline positive iPSC colonies formed per well) / (number of cells seeded before OKSM induction per well). D. The observed degree of overlap in clones forming iPSCs in each split for n = 3 independent biological replicates. E. Schematic of single-cell Rewind for retrospectively identifying hiF-T cells primed to reprogram into iPSCs. Here, we transduced hiF-T cells at an MOI of ∼0.15 with our barcode library. After 3 cell divisions, we sorted the successfully barcoded population (GFP positive) and divided the culture into two splits (A and B): with split A we reprogrammed the cells into iPSCs via induction of OKSM and performed barcode DNA-sequencing to identify primed clones and with split B (i.e., “carbon copy”) we immediately performed single-cell RNA-sequencing and barcode DNA-sequencing to label single-cell expression profiles with clone barcodes. F. We applied the Uniform Manifold Approximation and Projection (UMAP) algorithm within Seurat to the first 50 principal components to spatially visualize differences in gene expression in hiF-Ts before OKSM induction in split B (i.e., “carbon copy”). Left UMAP: Cells are colored by clusters determined using Seurat’s FindClusters command at a resolution of 0.45 (i.e., “Seurat clusters, res = 0.45”). Right UMAP: Cells are colored as primed (red) or nonprimed (light gray), determined by which clone twins formed iPSC colonies in reprogramming split A. Bar plot: We asked whether the primed hiF-T cells were more transcriptionally similar to each other than to the average of 1000 random samples of an equal number of nonprimed cells (i.e., “P(same distribution)”) and plotted the corresponding probability distributions across Seurat clusters (see Methods). G. We plotted the log2(fold change) between primed cells versus nonprimed cells from our data versus the log2(fold change) between hiF-T cells and hiF-T-derived iPSCs from (Cacchiarelli et al., 2015) for individual genes. Genes near the x-axis were differentially expressed between primed versus nonprimed cells but not between fibroblasts and iPSCs, while genes near the y-axis were vice versa. We labeled selected positive priming markers (in red) and negative priming markers (in blue). We chose 10 markers associated with iPSCs, fibroblasts, activated fibroblasts, and myofibroblasts identified in previous studies and asked if any broad category fit our observed priming markers. We plotted the average log2(fold change) between primed cells versus nonprimed cells for markers in each category and asked if the observed distribution was different from random (i.e., “P(median not random)”). H. Here, we transduced hiF-T cells at an MOI of ∼1 with our barcode library. After 3-4 cell divisions, we divided the culture into two splits (A and B): with split A we reprogrammed the cells into iPSCs via induction of OKSM and performed barcode DNA-sequencing to identify primed clones and with split B fixed cells *in situ* on glass slides before OKSM induction. We designed RNA clone barcode probes to label and identify primed clone barcodes by RNA FISH in the fixed cells (see Methods). We measured *OCT4* and *SOX2* expression in individual primed cells (in red) and nonprimed cells (in light gray) and plotted the population distribution of counts per cell for each gene. P-values comparing sample medians were calculated using the Wilcoxon rank sum test.

The application of Rewind requires that reprogrammability is largely dictated by heritable, intrinsic (meaning innate) differences in cellular state as opposed to extrinsic factors such as cell-cell interactions with neighbors or other microenvironmental factors. That way, the “carbon copy” cells would reflect the state of the cells that successfully reprogrammed. To demonstrate the primed state was intrinsic and heritable over at least a few cell divisions, we barcoded a population of 300,000-400,000 fibroblasts, let them proliferate for 3-5 divisions, and split each set of twins into different plates, hence randomizing their microenvironmental context. We then induced OKSM in both plates until the emergence of iPSCs (Figure 1B). If extrinsic factors were responsible for determining which cells reprogrammed following induction, we would expect largely distinct sets of barcodes in each population (maximum overlap of 0.385% across 10,000 simulations *in silico*) (Figure 1C). Instead, we observed a much higher degree of overlap (7.56%-37.1%) in barcodes even 14 days post-transduction, consistent with previous barcoding experiments (Shakiba et al., 2019; Yunusova et al., 2017) (Figures 1D, S2A-B). Thus, intrinsic differences that persist for at least several cell divisions are major determinants of cellular reprogramming, enabling us to apply Rewind in this system.

Pluripotent cells form colonies with large numbers of cells in them (>100 cells), hence we assumed that a large number of reads corresponding to a particular clone barcode was an indicator that those cells had successfully reprogrammed. To validate this assumption, we performed a similar barcode overlap experiment as before but with a split for generating embryoid bodies, which are cell aggregates mimicking the early embryo (Sheridan et al., 2012). We found that barcodes with a larger number of reads make up a majority of the resulting embryoid bodies, validating the use of number of reads as a proxy for pluripotency (Figure S2D). Furthermore, we found that primed twins even when separated form iPSC colonies of a similar size, as has been previously reported (Figure S2C) (Shakiba et al., 2019).

Having validated our ability to use Rewind in the hiF-T system, we applied it to measure gene expression differences between primed and nonprimed fibroblasts. We barcoded a population of fibroblasts as before in our barcode overlap experiments, let them divide for 3 divisions, and then separated the twins into different experimental splits. One split was reprogrammed into iPSCs via induction of the OKSM factors and the other split (i.e., “carbon copy”) was run through the single-cell RNA-sequencing pipeline, yielding gene expression profiles for individual cells. Because our clone barcodes are both integrated into the DNA and expressed into mRNA, clone barcodes can be both detected by targeted barcode DNA-sequencing and also detected in the single-cell transcriptomes generated using the 10x chromium platform (Goyal et al., 2021). By connecting both the clone barcode and the 10x cell ID (see Methods), we are able to assign the clone barcode to single-cell expression profiles in our dataset (Figure 1E). We filtered out all cells with spurious barcodes with unusual sequences or few reads and also removed all cells with multiple barcodes (Goyal et al., 2021). After those filtering steps, we were able to confidently label 28.9% of all cells as containing a single clone barcode. After sequencing the clone barcodes in the resulting iPSCs corresponding to primed cells, we additionally labeled cells as primed or nonprimed in the single-cell RNA-sequencing dataset before comparing gene expression differences. We identified 42 cells as primed in our whole population of 13,589 cells, consistent with what we would roughly expect based on our observed reprogramming efficiencies. To avoid biases resulting from barcoded and nonbarcoded cells (i.e., possible gene expression differences facilitating integration of lentiviral DNA or not), all comparisons between primed and nonprimed cells were done only with barcoded cells (Figure S3A).

### Primed cells in the initial population have measurable gene expression differences before OKSM exposure

Having identified the rare primed cells within our single-cell RNA-sequencing dataset, we could then compute the expression differences between primed and nonprimed cells to find transcriptome markers of the primed state (Figure 1F). Most of the markers we identified that distinguished primed and nonprimed cells were consistently differentially expressed across three biologically independent Rewind experiments (Figure S5A).

Upon categorizing the genes that were differentially expressed between primed and nonprimed fibroblasts, we found two general groupings. One consisted of genes involved in cell cycle regulation, pointing to an overall speedup of cell cycle progression in primed cells. Examples include increased expression of *MKI67, TOP2A*, and *CENPF*, all of which are expressed during the G2/M phase of the cell cycle. Increased expression of genes specific to the G2/M phase of the cell cycle may potentially indicate increased proliferation rate, because increases in proliferation rate are usually the result of a shortening of the G1 phase (Eastman & Guo, 2020), making expression of G2/M phase genes relatively higher within the population as a whole. As predicted, primed cells had a higher fraction of cells in G2/M (65.9% versus 32.3% in nonprimed) and a lower fraction of cells in G1 (9.8% versus 34.9% in nonprimed) (Figure S3B). Primed fibroblasts also had lower expression of cyclin dependent kinase inhibitors *CDKN2A* and *CDKN1A*, which are known reprogramming barriers (H. Li et al., 2009; Utikal et al., 2009; Zhan et al., 2019). Genes associated with M-phase regulation (*CENPF, SMC4, CDC20*) or microtubules (*ASPM, TUBA1A*) were also upregulated in primed fibroblasts.

The other primary gene expression signature of primed fibroblasts we observed was lower expression of several genes associated with activated fibroblasts (*SPP1, SERPINE2, THBS1, TAGLN*) (Hsia et al., 2016; Layton et al., 2020; Peyser et al., 2019; Sandberg et al., 2019), differentiation into myofibroblasts (*MYL9, ACTA2, TPM2, POSTN*) (Guerrero-Juarez et al., 2019; Hsia et al., 2016; Layton et al., 2020; Walker et al., 2019), and pathological fibrosis (*GDF15, IGFBP7, GAS5*) (L.-X. Liu et al., 2009; Radwanska et al., 2022; Tang et al., 2020)(Radwanska et al., 2022; Tang et al., 2020). This signature suggests that unactivated fibroblasts within the population are more likely to reprogram. We wondered if these priming markers might be regulated by a core set of common transcription factors. We used a database that aggregates lists of regulatory relationships between transcription factors and target genes to identify potential common transcription factors for the positive and negative priming markers separately (Keenan et al., 2019). Interestingly, we found that our negative priming markers were positively associated with several transcription factors known to either switch on during EMT or off during MET, including *TWIST2, SNAI2, OSR1*, and *PRRX2* (Figures S4C-D) (Mellis et al., 2021). We manually identified a number of binding motifs for these factors upstream of *SPP1* and *FTH1* (Figure S4E).

We wanted to confirm differential expression of the positive and negative priming markers by direct visualization of gene expression by single-molecule RNA FISH. We performed Rewind experiments similar to those in Figure 1E, but instead of subjecting one split of the experiment to single-cell RNA-sequencing, we instead immediately fixed the “carbon copy” fibroblasts after splitting. After identifying barcodes corresponding to iPSCs in the split in which OKSM was induced, we designed RNA FISH probes targeting the clone barcodes of primed cells (Figure 2A). In our fixed samples, we identified primed cells using these clone barcode RNA FISH probes (Figure S5C). We further measured expression of positive priming markers *TOP2A* and *CENPF* as well as negative priming markers *SPP1* and *SQSTM1*, confirming that they had higher expression and lower expression respectively in primed cells as compared to nonprimed cells (Figure 2B). Single-molecule RNA FISH also confirmed that there was no difference in *OCT4* or *SOX2* expression between primed and nonprimed cells, in line with our single-cell RNA-sequencing results and eliminating the possibility that leaky expression of OKSM before induction could be responsible for priming individual cells for reprogramming (Figure 1H).

**Figure 2:**
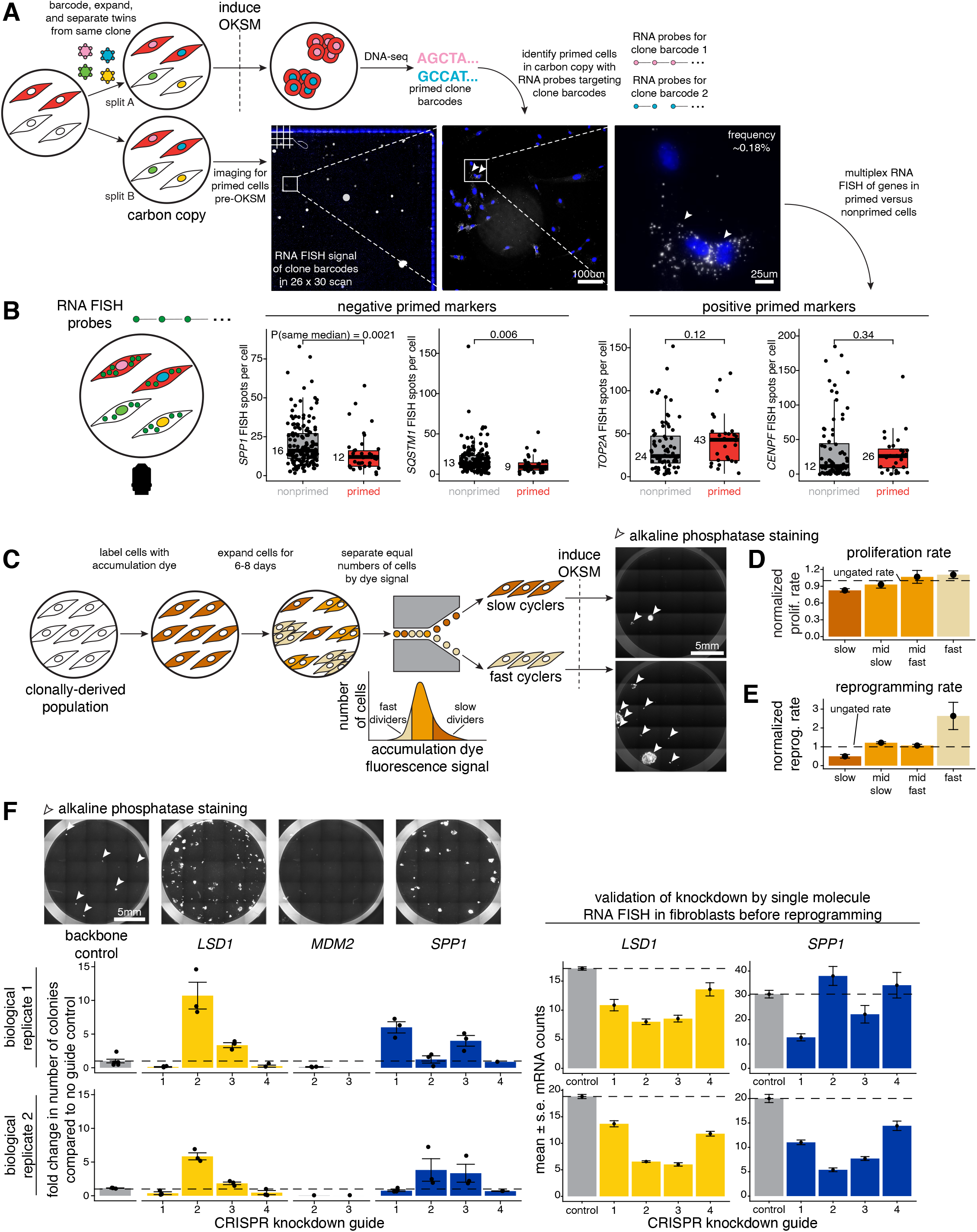
Selecting for positive and negative priming markers in the initial population predictably changes reprogramming outcome. A. Schematic of *in situ* Rewind for retrospectively identifying hiF-T cells primed to reprogram into iPSCs fixed on slides. Here, we transduced hiF-T cells at an MOI of ∼1 with our barcode library. After 3-4 cell divisions, we divided the culture into two splits (A and B). With split A we reprogrammed the cells into iPSCs via induction of OKSM and performed barcode DNA-sequencing to identify primed clones. With split B (i.e., “carbon copy”), we immediately fixed cells on slides after splitting but before OKSM induction. We marked primed cells in split B with RNA FISH probes to primed clone barcodes identified in the reprogrammed iPSCs in split A. We imaged DAPI (blue) and RNA FISH signal from primed clonal barcodes (white) at different magnifications on a fluorescence microscope. B. We measured gene expression in individual primed and nonprimed cells in split B by performing single-molecule RNA FISH for *SPP1* and *SQSTM1* (i.e., negative priming markers) as well as *TOP2A* and *CENPF* (i.e., positive priming markers). For each marker, we plotted FISH spots per cell for many primed (in red) and nonprimed (in gray) cells. P-values comparing sample medians were calculated via Wilcoxon rank sum test. C. To sort cells based on cycling speed, we stained cells with a fluorescent dye that becomes diluted with successive cell divisions (i.e., “accumulation dye”). Here, lighter shades of brown indicate higher dilution and more divisions (i.e., “fast cyclers”) while darker shades of brown indicate lower dilution and fewer divisions (i.e., “slow cyclers”). We sorted out slow and fast cells, reprogrammed each into iPSCs via induction of OKSM, and quantified the number of alkaline phosphatase-positive iPSC colonies (in white) formed. D. We sorted cells based on cycling speed into four bins: slow, mid slow, mid fast, and fast. We plated cells, let them divide for 2-3 days, and measured the proliferation rate, where proliferation rate = (number of cells at the end) / (number of cells seeded). All measured proliferation rates were normalized to a control of ungated hiF-T cells (i.e., “normalized prolif. rate”). Metric shown is mean +/- standard error for n = 4 independent biological replicates for slow and fast and n = 2 for mid slow and mid fast. E. We sorted cells based on cycling speed into four bins: slow, mid slow, mid fast, and fast. We reprogrammed each bin into iPSCs via induction of OKSM and quantified the number of iPSC colonies formed. The number of iPSC colonies formed for each bin was normalized to a control of ungated hiF-T cells (i.e., “normalized reprog. rate”). Metric shown is mean +/- standard error for n = 4 independent biological replicates and n = 3 for mid slow and mid fast. F. We designed CRISPR guides to knockdown mRNA expression of *LSD1* (positive control, in yellow), *MDM2* (negative control, in gray), and *SPP1* (in blue). For each gene knockdown condition we chose a representative image to visualize the number of alkaline phosphatase-positive iPSCs formed upon reprogramming. We quantified the number of iPSC colonies formed for each guide (labeled numerically) for each gene and reported each measurement as a fold change value in comparison to a no guide control (i.e., backbone vector lacking a targeting guide RNA). For each CRISPR guide knocking down *LSD1* or *SPP1*, we measured the expression level of each respective gene with single-molecule RNA FISH. Metric shown is mean +/- standard error values from measurements from many individual cells.

The transcriptional profile of primed cells could indicate whether the primed state is directed towards a target fate or not (i.e., “head start priming”). In head start priming, a cell primed towards an iPSC could express some iPSC-specific genes, giving those cells a “head start” towards reprogramming (Nemajerova et al., 2012; Wakao et al., 2011). To look for head start priming, we compared the transcriptional profile of priming from our Rewind data to existing bulk RNA-sequencing data across multiple time points during iPSC reprogramming (after induction of reprogramming) in our cell line (Cacchiarelli et al., 2015). Broadly, positive priming markers increased in expression through the reprogramming process while negative priming markers decreased in expression early on in reprogramming (Figures S4A-B). The positive priming markers, however, did not show much increased expression in the reprogrammed iPSCs (Figure 1G). The negative priming markers showed a more significant downregulation as cells reprogrammed from fibroblasts to iPSCs, showing that the loss of expression of these (primarily fibroblast-specific) genes continued during the fibroblast to iPSCs transition.

Furthermore, we looked at the expression of sets of genes specific to iPSCs, fibroblasts, activated fibroblasts, and myofibroblasts in primed versus nonprimed cells (Figure 1G). The expression of canonical iPSC genes (e.g., *NANOG, LIN28A, DNMT3B*) was virtually unchanged in primed vs. nonprimed cells, while many activated fibroblast and myofibroblast genes were significantly downregulated in primed cells. These results argue against primed cells showing any indication of a head start towards the iPSC fate, instead suggesting an association with a generic lack of fibroblast activation.

### Selecting for positive and negative priming markers in the initial population predictably changes reprogramming outcome

While the transcriptional signatures of primed cells suggested the importance of cell cycle speed and fibroblast activation (here, meaning expression of markers of fibroblast activation or myofibroblast differentiation) in reprogrammability, we wanted to confirm the associations between cell cycle and lack of fibroblast activation with priming via alternative methods. In the case of cell cycle, several studies have shown that cells already undergoing the reprogramming process can show increased rates of division (Babos et al., 2019; S. Guo et al., 2014; Smith et al., 2010), but less is known (Utikal et al., 2009) about how natural variability in proliferation rate *before* the reprogramming process is associated with reprogrammability.

To measure how cell cycling speed in uninduced fibroblasts affected the ability of cells to reprogram, we separated cells by cycling speed by staining them with a fluorescent dye that becomes diluted with successive cell divisions (Figure 2C). After a sufficient number of divisions, fast cycling cells were identified with low fluorescent signal while slow cycling cells were identified with high fluorescent signal due to differential dye diffusion. Upon sorting, we found that fast cycling cells proliferated at a 1.10-fold faster rate compared to the ungated control and a 1.34-fold faster rate compared to slow cycling cells (Figure 2D). Upon induction of OKSM in these different subpopulations, we found that fast cycling cells generated 2.64-fold more colonies than ungated cells and 5.32-fold more colonies than slow cycling cells (Figure 2E). There has been a report of an ultra-fast cycling (8-fold faster) population with higher reprogramming potential (S. Guo et al., 2014); we did not observe such a subpopulation in our cell line.

Why do faster cycling cells correlate with more reprogramming into iPSCs? The difference could merely be the result of increased numbers of cells prior to induction owing to the increased number of divisions in fast cycling cells, or it could be that these cells have an intrinsically higher propensity to reprogram. To directly measure if primed clones reprogrammed more efficiently solely due to entering the reprogramming process with more cells, we measured the number of cells per clone for primed and nonprimed cells at the time of OKSM induction in our Rewind single-cell RNA-sequencing dataset from Figure 1E (Figure S6D). We observed a minimal difference in the distribution number of starting cells for primed versus nonprimed cells (mean(primed) = 1.20 versus mean(nonprimed) = 1.25, p-value = 0.57), indicating that reprogramming success is not merely a function of number of progenitors before OKSM induction.

Given that the cells were only kept in culture for 2-3 divisions after barcoding but before splitting for single-cell RNA-sequencing, we wondered how sensitive this approach was in detecting expected differences in number of starting cells. We performed an additional Rewind experiment in which we sorted our “carbon copy” split into different groups based on cycling speed before performing single-cell RNA-sequencing. With those data, we measured the starting number of cells per clone across cycling speeds as an indicator of proliferation rate.

While we were able to detect differences in the number of starting cells between fast and ungated cells, we did not detect a difference between slow and ungated cells. We cannot distinguish whether the minimal difference in number of starting cells between slow and ungated cells here is because of smaller differences in cycling speed between these groups in this experiment or because of the inherent noisiness in measuring number of starting cells per clone in our dataset. Despite this limitation, we saw minimal differences in distribution of number of starting cells between primed and nonprimed cells both in bulk and when separating cells into cycling speed sort groupings. We also demonstrated that a higher fraction of fast clones reprogram into iPSCs compared to ungated and slow clones (Figure S6B). Additionally, we estimated that fast clones and slow clones would need to maintain their respective pre-OKSM cycling speeds for nearly a week following OKSM induction, by which point reprogramming is well underway, to generate a sufficiently different number of progenitors to fully explain the different observed reprogramming rates (Figure S6E). These results collectively argue that naturally-occurring fast cycling cells within an otherwise homogenous population have an intrinsically higher rate of reprogramming than slow cycling cells.

While the above results show that cycling speed is associated with higher rates of reprogramming, not every fast cycling cell underwent reprogramming. Our molecular profiling results suggested that lower levels of fibroblast activation markers (Figure 1G) may be another important feature of priming. To demonstrate that a lack of fibroblast activation markers *per se* could lead to increase of reprogramming efficiency, we knocked down the expression of selected negative priming markers, including some with potential roles in fibroblast activation, to see if their loss could lead to an increased number of primed cells. We used CRISPR/Cas9 to knock down the negative priming markers *SPP1, FTH1*, and *CDKN1A*, as well as putative upstream regulators *MYBL2* and *NFE2L2* identified by motif analysis (Figure S5B). We included *LSD1* and *MDM2* knockdowns as positive and negative controls: *LSD1* knockdown is known to increase reprogramming rate, and *MDM2* knockdown is known to block reprogramming via upregulating p53 activity (Cacchiarelli et al., 2015; Wienken et al., 2016). We focused on *SPP1* given its known role in mediating TGF-B induced fibroblast activation (Kramerova et al., 2019; Lenga et al., 2008). Across two biologically independent experiments, knockdown of *SPP1* mRNA expression resulted in a higher reprogramming efficiency compared to cells infected with the same CRISPR lentivirus without a guide RNA (Figure 2F). Different guides knocked down RNA to different extents (including variability across replicates); we found that the greater the level of mRNA knockdown, the higher the number of iPSC colonies formed was. Notably, we did not observe as clear of an association between knockdown of SPP1 at the protein level and iPSC colony formation rate; however, this lack of association could be explained by nonspecific SPP1 antibody binding in our assay or be due to SPP1 being a secreted protein (Figure S5D) (Rittling & Feng, 1998). These results show that knocking down factors that may drive fibroblast activation can directly increase the ability of fibroblasts to reprogram into iPSCs; notably, this knockdown did not affect cell cycle speed (Figure S5E).

### Cycling speed and fibroblast activation are aspects of a single axis of biological variability marking primed cells

Given that we identified two modes of priming (faster cycling and lack of fibroblast activation), we wondered the extent to which these two modes were distinct, or whether they were just different aspects of a single underlying axis of biological variability. To distinguish these possibilities, we performed a Rewind experiment as before in Figure 1E; however, with our “carbon copy” split we sorted out cells based on cycling speed and then ran each group separately through the single-cell RNA-sequencing pipeline (Figures 3A-B). Simultaneously quantifying cycling speed and priming marker expression in individual cells enabled us to directly measure the relative contribution of each mode of priming to overall priming status.

**Figure 3:**
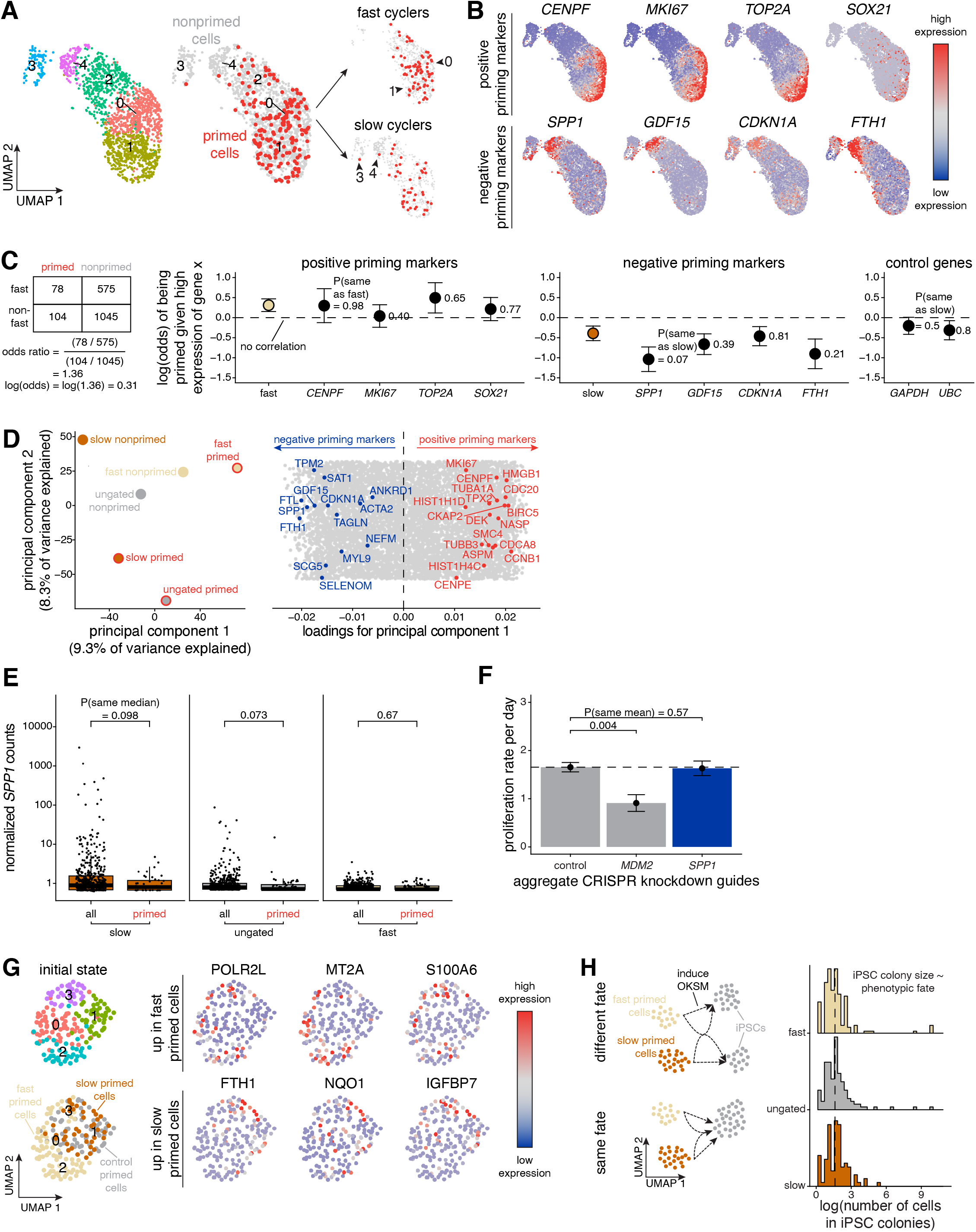
Cycling speed and fibroblast activation are aspects of a single axis of biological variability marking primed cells. A. Here, we simultaneously stained hiF-T cells with an accumulation dye and transduced the same hiF-T cells at an MOI of ∼0.15 with our barcode library. After 2-3 cell divisions, we divided the culture into two splits. With one split, we reprogrammed the cells into iPSCs via induction of OKSM and performed barcode B. DNA-sequencing to identify primed cones. With the other split, we sorted out successfully barcoded cells (GFP positive) and sorted out equal numbers of slow, ungated, and fast cells based on the accumulation dye signal. We immediately performed single-cell RNA-sequencing on each separate cycling speed population and barcode DNA-sequencing to label single-cell expression profiles with clone barcodes. We applied the Uniform Manifold Approximation and Projection (UMAP) algorithm to the first 50 principal components to visualize differences in gene expression in hiF-Ts before OKSM induction. Shown are cells for which we could confidently assign a single clone barcode across all cycling speeds on the left UMAPs and in fast cyclers versus slow cyclers on the right UMAPs. Cells are colored by clusters determined using Seurat’s FindClusters command at a resolution of 0.3 or by priming status (primed in red, nonprimed in gray). On the UMAP, we recolored each cell for its expression for a select subset of positive and negative priming markers using Seurat’s FeaturePlot command. High expression is marked in red while low expression is marked in blue. Positive priming markers *CENPF, MKI67, TOP2A*, and *SOX21* are primarily expressed in clusters 0 and 1. Negative priming markers *SPP1, GDF15, CDKN1A*, and *FTH1* are primarily expressed in clusters 3 and 4. C. To measure the relative explanatory power of cycling speed and a subset of our identified priming markers (as well as the housekeeping genes *GAPDH* and *UBC*) in the context of priming, we calculated and plotted odds ratios (see Methods). The odds ratios for each gene was calculated with corresponding standard error separately in n = 3 biologically independent single-cell RNA-sequencing datasets and aggregated via a random-effects model using the metafor package in R. D. To visualize different axes of biological variability in our single-cell RNA-sequencing dataset, we aggregated expression profiles in each of our cycling speed-priming categories using Seurat’s AggregateExpression command and plotted the aggregates in principal component space. We extracted loadings for each principal component and plotted them for principal component 1, highlighting the positive priming markers (in red) and negative priming markers (in blue). E. To evaluate the correlation between *SPP1* levels and cycling speed, we plotted normalized SPP1 counts from our single-cell RNA-sequencing dataset described in Figure 3A for all cells versus primed cells within each cycling speed. Each dot represents the normalized *SPP1* counts from an individual cell. F. Here, we plotted aggregated proliferation rates for all guides to a given gene target. Metric shown is mean +/- standard error for all aggregated guides. The proliferation rates for individual guides across n = 2 independent biological replicates and details on how we calculated proliferation rate can be found in Figure S5E. P-values comparing sample means were calculated using the Student’s t-test. G. We subsetted our single-cell RNA-sequencing dataset described in Figure 3A to include only primed cells and clustered using the Uniform Manifold Approximation and Projection (UMAP) algorithm to the first 50 principal components to visualize difference in gene expression between slow primed, ungated primed, and fast primed cells. On the UMAP, we recolored each cell for its expression for a subset of markers that were differentially expressed between fast primed cells and slow/ungated primed cells. High expression is marked in red while low expression is marked in blue. H. To determine if the iPSC colonies arising from fast versus slow cycling primed cells had any phenotypic differences, we plotted the distribution of iPSC colony size for iPSCs derived from slow, ungated, and fast clones. The size of each iPSC colony was determined by normalizing read counts after performing DNA-sequencing on the reprogrammed iPSCs using spike-ins of known cell number and of known barcodes.

We first set about determining their relative contributions by using odds ratios between reprogrammability as a function of proliferation speed and transcriptional profile. For example, we asked what are the odds that a fast cycling cell is also primed? We compared odds ratios for cycling compared to several positive (*CENPF, MKI67, TOP2A, SOX21*) and negative (*SPP1, GDF15, CDKN1A, FTH1*) priming markers. The positive priming markers, which were predominantly associated with cell cycle progression, did not have a stronger positive association with priming compared with fast cycling. However, several of the negative priming markers, in particular *SPP1*, had a stronger negative association with priming compared with slow cycling (Figure 3C). To use information lost by dichotomizing continuous gene expression values to calculate odds ratios, we generated logistic regression models in which we determined the contributions of cycling speed, expression of each gene, and the interaction between these terms in predicting priming. Again, we saw that the negative priming markers explained more of the variation in priming status compared to cycling speed (Figure S7A).

We wondered if this difference in explanatory power was because the negative priming markers, associated with fibroblast activation, represented a distinct axis of priming. To answer this question, we first aggregated molecular profiles from cells in each of our cycling speed and priming categories and visualized the similarity in principal component space (Figure 3D). The aggregates separated by cycling speed along principal component 1, which was determined primarily by our identified priming markers (i.e., positive priming markers in one direction and negative priming markers in the other direction) and argued for a single shared axis of variability. Notably, slow primed and ungated primed cells separated from nonprimed cells along principal component 2, which explained a similar amount of variation in our aggregate samples yet did not correlate with our identified priming markers. This principal component hints at additional, unidentified axes of biological variability driving priming distinct from the axis we have described in detail here.

To evaluate the relationship between cycling speed and fibroblast activation more closely, we focused on *SPP1* because it was the negative marker with the highest explanatory power and earlier we had shown how expression levels of *SPP1* can directly affect reprogramming efficiency. When we measured *SPP1* mRNA levels between primed and nonprimed cells across cycling speeds, *SPP1* expression was strongly anti-correlated with cycling speed broadly but was generally lower among primed cells compared to nonprimed cells within each cycling speed group (Figure 3E). This pattern of *SPP1* expression could indicate that slow cycling primed cells may successfully reprogram *in spite* of having a slow cycling speed (in part by having low *SPP1* levels) or that slow cycling primed cells are more simply marked by a relatively faster cycling speed compared to their nonprimed peers. To distinguish these possibilities, we measured the fraction of cells in G1 as a proxy for cycling speed (i.e., a higher fraction of cells in G1 implies a slower cycling speed) (Eastman & Guo, 2020) across cycling speed categories and between primed and nonprimed cells. We found that across all cycling speed categories, primed cells had a lower fraction of cells in G1; surprisingly slow primed cells had an even lower fraction of cells in G1 compared with fast cyclers in bulk. Additionally, cells with the lowest 2.5% and 5% of *SPP1* expression had a significantly lower fraction of cells in G1 compared to the remaining population (Figure S6F). However, slow primed cells demonstrated fewer (0.28 cells fewer on average, p-value = 0.16) mean starting cells per clone compared to fast primed cells. As such, we cannot conclude whether slow primed cells are misclassified as slow by our accumulation dye approach or somehow spend relatively less time in G1 yet still take a longer overall time to progress through the entire cell cycle.

Given the correlation between *SPP1* mRNA levels and cycling speed, we wondered if knockdown of *SPP1* may increase reprogramming efficiency by increasing cycling speed. When we knocked down *SPP1* mRNA levels by CRISPR in Figure 2F, we found that cycling speed was unchanged in cells with guides to *SPP1* compared to control (Figure 3F). In contrast, knockdown of *MDM2* mRNA levels significantly reduced cycling speed and reprogramming efficiency, presumably via upregulation of p53 activity. That *SPP1* knockdown can seemingly increase reprogramming rate independently of cycling speed indicates that fast cycling speed, per se, is not required for successful iPSC reprogramming.

These results collectively show that fast cycling and lack of fibroblast activation genes likely mark a single underlying axis of variability, meaning that fast cycling cells show low levels of fibroblast activation markers and vice versa. While reducing *SPP1* levels can increase reprogramming efficiency without changing cycling speed, we do not know if the opposite is also true; indeed, cells with fast cycling speed may only affect reprogramming by virtue of their association with lack of fibroblast activation. Of these modes of priming, however, lack of fibroblast activation markers was a stronger predictor of whether an individual cell was likely to reprogram.

In principal component space, fast cycling cells seemed to differ from the slow cycling and ungated cells particularly in the second principal component. When we clustered solely the primed cells (as opposed to the entire population) and projected into UMAP space, we indeed found that fast cycling primed cells occupied different clusters than the ungated and slow primed cells (Figure 3G). Thus, fast cycling primed cells may represent a cellular state distinct from that of slow cycling primed cells, showing that there are multiple types of primed cells. A parallel question is whether the iPSC colonies arising from fast versus slow cycling primed cells had any phenotypic differences; we found that colonies arising from both types of cells did not have any appreciable differences in the number of cells per colony (Figure 3H) (although when only large colonies are considered, fast primed cells did lead to larger iPSC colonies; Figure S7G). These results suggest that different initial states may adopt similar ultimate fates as part of the reprogramming process.

### The primed state is defined by extrinsic perturbations in addition to intrinsic cell state

Here, we have defined primed cells as cells that are able to undergo reprogramming when induced. We have demonstrated a mapping between the intrinsic state of the cell and reprogramming outcome, seemingly enabling a purely intrinsic, state-based definition of priming. There are perturbations, however, that can change the apparent efficiency of reprogramming, challenging that assertion.

One such perturbation is inhibition of LSD1 (histone lysine demethylase 1), which was recently identified as a reprogramming booster in our specific cell line (Cacchiarelli et al., 2015) and for which a chemical inhibitor is readily available. If priming were purely intrinsic, then the apparent boost in reprogramming would have to come from an increase in the proliferation of those same existing intrinsically primed cells. If, on the other hand, LSD1 inhibition allowed cells to reprogram that otherwise would not have reprogrammed, then one could say that the perturbation acts by a “reclassification of state” for priming, consequently meaning that priming cannot be defined purely intrinsically (Figure 4A). To discriminate between these possibilities, We barcoded fibroblasts and separated twins into two splits, one with pure OKSM induction (i.e., “control”) and the other with both LSD1 inhibition and OKSM induction (i.e., “+LSD1i”). Upon sequencing the resultant iPSC colonies for clone barcodes, we found that a number of clone barcodes showed up in both the pure OKSM and the LSD1 inhibition conditions, but an even larger proportion showed up only when LSD1 was inhibited (Figure 4B). Cells primed for reprogramming only with LSD1 inhibition still exhibited memory for the primed state to roughly the same extent as conventional priming (Figure 4C). Thus, LSD1 inhibition concurrent with OKSM induction led to a reclassification of initial cell states as primed.

**Figure 4:**
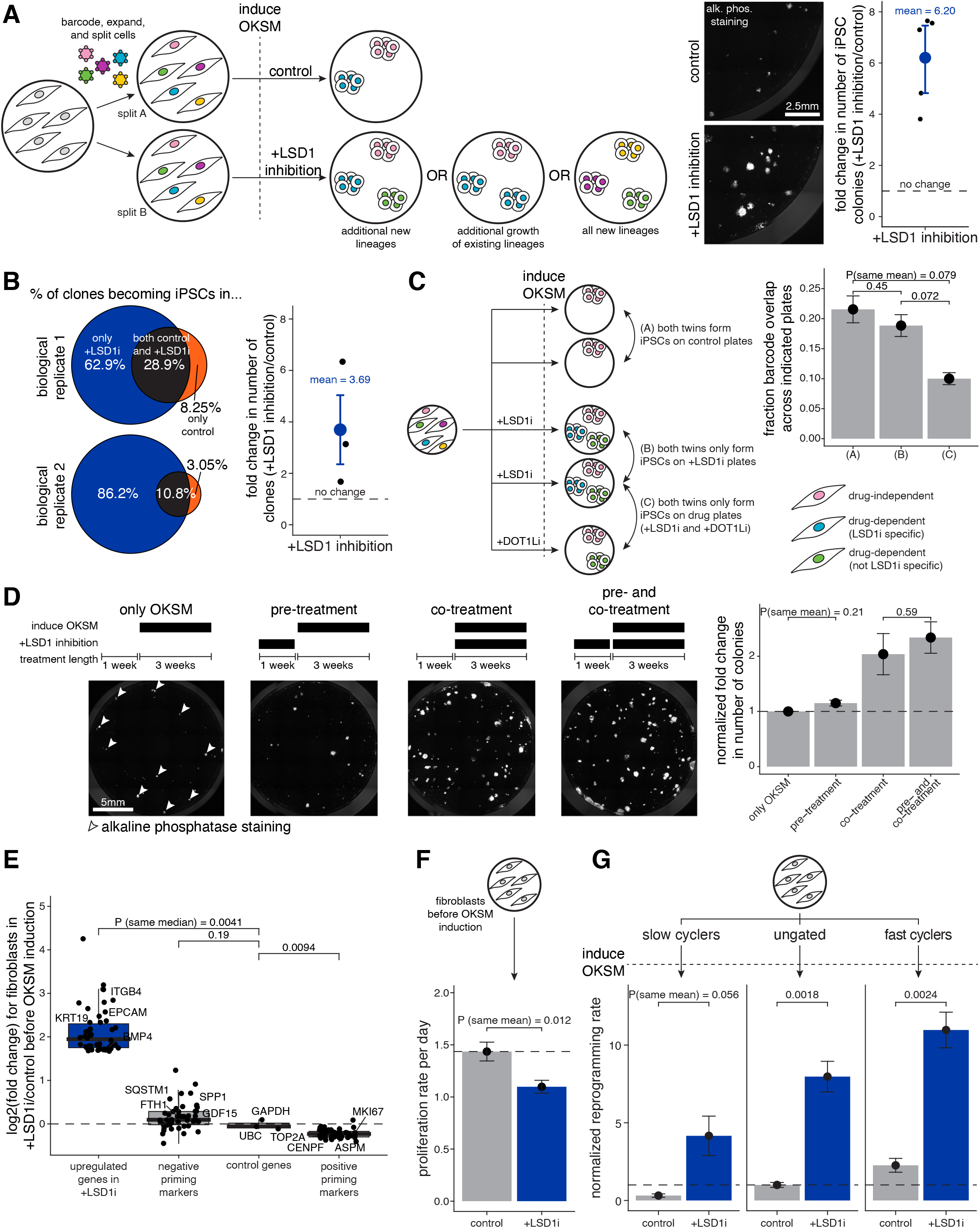
The primed state is defined by extrinsic perturbations in addition to intrinsic cell state. A. Schematic of Rewind for following fates of hiF-T cells reprogrammed in OKSM alone (i.e., “control”) and in OKSM with LSD1 inhibition (i.e., “+LSD1 inhibition”) with different possibilities for the source of additional colonies described. We transduced hiF-T cells at an MOI of ∼1 with our barcode library. After 3-4 cell divisions, we divided the cultures into splits (A and B). In split A we reprogrammed with OKSM alone while in split B we reprogrammed with OKSM and LSD1 inhibition. After reprogramming, we stained for alkaline phosphatase activity (in white) and imaged using fluorescence microscopy. We measured the number of colonies formed in each reprogramming condition and plotted the fold change between OKSM with LSD1 inhibition over OKSM alone. Metric shown is mean +/- standard error for n = 5 independent biological replicates. B. For some experiments described in Figure 4A, we performed barcode DNA-sequencing on iPSCs formed in OKSM alone (i.e., “control”) versus in OKSM with LSD1 inhibition (i.e., “+LSD1i”). We compared barcodes across each reprogramming condition after performing normalizations described in Figure S8. We measured the number of clones in the reprogrammed iPSCs in each reprogramming condition and plotted the fold change between OKSM with LSD1 inhibition over OKSM alone. Metric shown is mean +/- standard error for n = 3 independent biological replicates. C. We performed a similar experiment as described in Figure 4A, but after 5-6 cell divisions we divided the cultures into five splits: two splits were reprogrammed with OKSM alone, two splits were reprogrammed with OKSM and LSD1 inhibition (i.e., “+LSD1i”), and one split was reprogrammed with OKSM and DOT1L inhibition (i.e., “+DOT1Li”). After reprogramming, we performed barcode DNA-sequencing on iPSCs formed in each reprogramming condition and compared barcode overlap within and across conditions as indicated. We plotted barcode overlap across each indicated comparison. Metric shown is mean +/- standard error for n = 2 independent biological replicates. P-values comparing sample means were calculated using the Student’s t-test. D. To determine when LSD1 inhibition acts to increase the number of iPSC colonies, we reprogrammed hiF-T cells via OKSM induction and added LSD1 inhibitor during the different time frames indicated. Shown are representative images of 24-wells after reprogramming and staining for alkaline phosphatase activity (in white) in each condition. We quantified and plotted the number of colonies in each condition after normalizing to the number of colonies formed in baseline reprogramming (i.e., “only OKSM”). Metric shown is mean +/- standard error for n = 2 independent biological replicates. P-values comparing sample means were calculated using the Student’s t-test. E. We performed bulk RNA-sequencing on hiF-T cells after one week in normal culture conditions versus culturing with LSD1 inhibition (i.e., “+LSD1i”) and performed differential expression analysis in DESeq2. OKSM was not induced in either condition. We plotted log2(fold change) values for different categories of genes in LSD1 inhibition over control. Shown from left to right are genes with the top 50 log2(fold change) values in the differential expression analysis (i.e., “upregulated genes in +LSD1i”), our top 25 negative priming markers, four housekeeping genes (*UBC, GAPDH, PGH1, ACTB*), and our top 25 positive priming markers. P-values comparing sample medians were calculated using the Wilcoxon rank sum test. F. To determine if LSD1 inhibition had any effect on proliferation rate, we measured proliferation rate per day as described in Figure S5E in hiF-T cells after one week in normal culture conditions (i.e., “control”) versus culturing with LSD1 inhibition (i.e., “LSD1i”). Metric shown is mean +/- standard error for n = 2 independent biological replicates. P-values comparing sample means are calculated using the Student’s t-test. G. We used the accumulation dye approach in Figure 2C to sort out hiF-Ts by cycling speed (slow, unsorted, and fast) and then reprogrammed each population with OKSM alone versus OKSM and LSD1 inhibition. After reprogramming, we stained the iPSCs with alkaline phosphatase and counted the number of iPSC colonies formed in each condition. We calculated the normalized reprogramming rate by dividing the number of colonies formed in each condition by the number of colonies formed in the unsorted population reprogrammed with OKSM alone. Metric shown is mean +/- standard error for n = 2 independent biological replicates. P-values comparing means were calculated using the Student’s t-test.

LSD1 inhibition *before* OKSM induction might also function to increase iPSC reprogramming efficiency by pushing nonprimed cells from the population into the same intrinsic state that was classified as primed without LSD1 inhibition. To test this possibility, we used inhibition of LSD1 at various points before and during reprogramming. We found that a 7 day pre-treatment with LSD1 inhibitor prior to OKSM induction did not lead to any appreciable difference in the number of iPSC colonies (Figure 4D), suggesting that LSD1 inhibition primarily changes the probability of a cell in a given state to reprogram rather than changing the state to resemble a specific primed state itself.

Supporting this conclusion, we found that LSD1 inhibition did not change the expression of our previously identified priming markers when comparing bulk RNA-sequencing profiles for hiF-Ts after a week in culture with or without LSD1 inhibition (Figure 4E). We did, however, observe upregulation of epithelial markers (*KRT19, EPCAM*) as well as factors whose expression early on following OKSM induction is associated with reprogramming success (*BMP4, ITGB4*) (L. Guo et al., 2019; Hayashi et al., 2016), consistent with LSD1’s proposed role in facilitating MET (Cacchiarelli et al., 2015) (Figure 4E). Additionally, knockdown of LSD1 by chemical inhibition or CRISPR did not increase and perhaps decreased cell cycling (Figure 4F) (Cacchiarelli et al., 2015; Sun et al., 2016), and LSD1 inhibition increased iPSC generation efficiency regardless of cycling status in uninduced fibroblasts (Figure 4G). Thus, LSD1 inhibition does not increase priming by pushing cells into a state of faster cycling. Together, these results are consistent with a model in which LSD1 inhibition allows cells to reprogram that would not have reprogrammed otherwise, and points to the fact that the primed cell state is not a single discrete state per se, but rather is a set of states whose extent is determined by the nature of the reprogramming induction itself.

We further wondered if cells that required LSD1 inhibition to reprogram also were able to reprogram when treated with other reprogramming boosters. Inhibition of DOT1L, an H3K79 methyltransferase, is known to also facilitate iPSC reprogramming (Onder et al., 2012; Wille & Sridharan, 2022). We found that some clones dependent on LSD1 inhibition could also form iPSCs with DOT1L inhibition, but the amount of memory across perturbation conditions was lower, implying a combination of booster specific and booster general mechanisms for expanding the subset of primed cells (Figure 4C).

### Clone of origin has less influence than reprogramming conditions in dictating the final molecular states of cells subjected to iPSC reprogramming

Above, we showed that LSD1 inhibition allows some cells to reprogram that would normally not reprogram. This result raises the question of whether the different initial state of these cells propagates to differences in the molecular profile of the ultimate iPSCs formed. Indeed, more generally, we wondered to what extent either the cell of origin or reprogramming conditions affected the ultimate iPSC state.

Given the heterogeneity in reprogrammed iPSCs (Nguyen et al., 2018; Xing et al., 2020; Yang et al., 2021), we needed to have single-cell resolution of these outcomes as well as information about the clonal origin of iPSC colonies. To trace reprogramming outcomes from the originating cell through to the final state with single-cell resolution, we used a method called FateMap (Goyal et al., 2021). In FateMap, we again barcode cells before inducing OKSM but we collect the entire pool of cells after reprogramming is finished instead of beforehand and perform single-cell RNA-sequencing on them. Thus, we can measure the transcriptional heterogeneity within and between individual iPSC clones. As reported, there was significant heterogeneity within the reprogrammed population as a whole (Figure 5A). Upon clustering, we found several distinct iPSC clusters, identifiable by broad average expression of pluripotency markers (clusters 0, 1, 2, 3). Other clusters included surviving fibroblasts (cluster 4) and a more indeterminate cluster potentially representing incomplete reprogramming or differentiating iPSCs (cluster 5) (Figures 5A-B).

**Figure 5:**
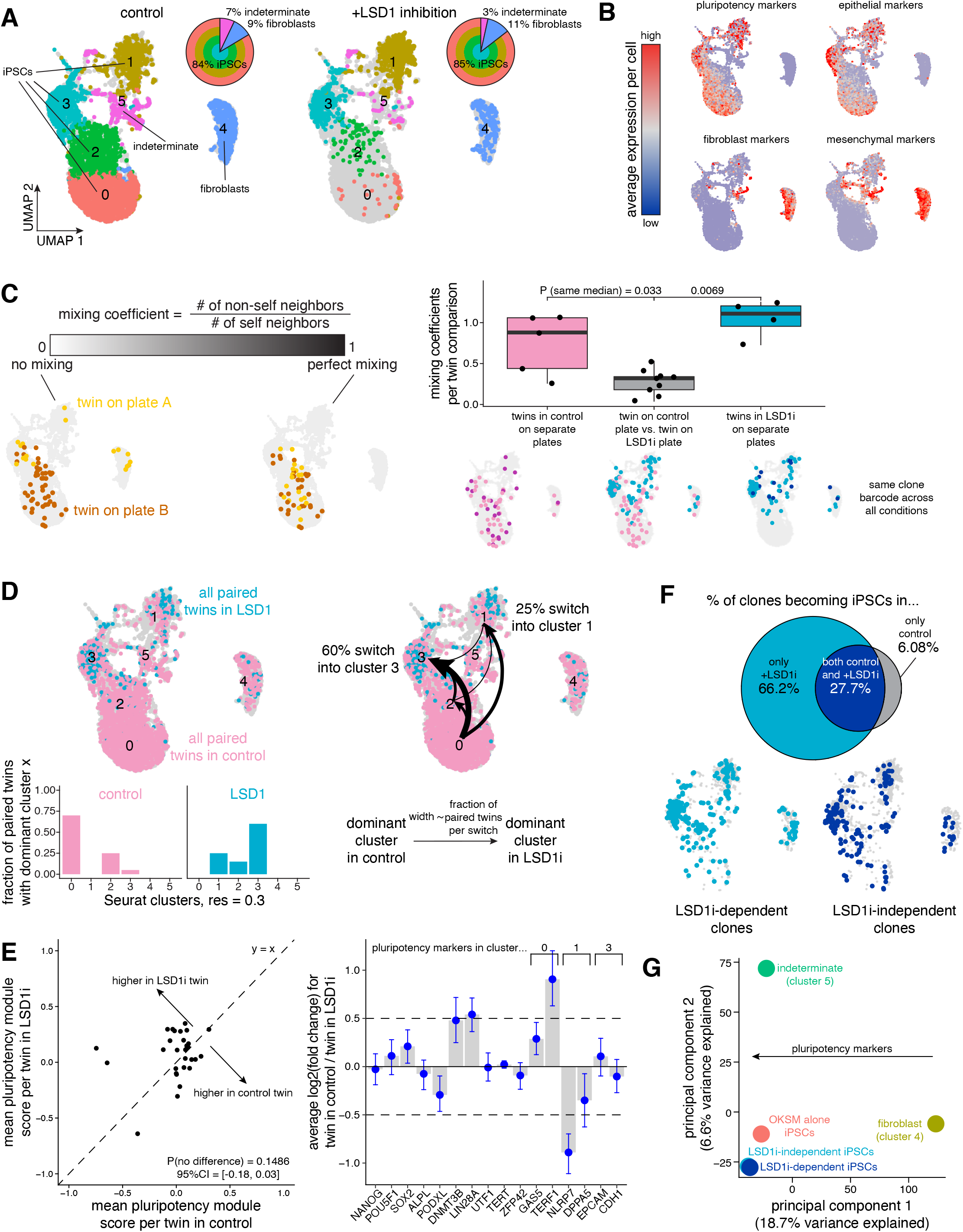
Clone of origin has less influence than reprogramming conditions in dictating the final molecular states of cells subjected to iPSC reprogramming. A. We transduced hiF-T cells at an MOI of ∼0.15 with our barcode library. After 3 cell divisions, we sorted the successfully barcoded population (GFP positive) and divided the culture into four splits for reprogramming: two splits reprogrammed with OKSM alone and two splits reprogrammed with OKSM and LSD1 inhibition. After reprogramming each split into iPSCs, we performed single-cell RNA-sequencing and barcode DNA-sequencing to label single-cell expression profiles with clone barcodes. We applied the Uniform Manifold Approximation and Projection (UMAP) to the first 50 principal components to spatially visualize differences in gene expression in the resulting iPSCs within and across each reprogramming condition. Cells are colored by cluster determined by using Seurat’s FindClusters command at a resolution of 0.3. For each reprogramming condition, we calculated the fraction of cells across iPSCs (clusters 0, 1, 2, 3), an indeterminate subset somewhere between iPSCs and fibroblasts (cluster 5, “indeterminate”), and fibroblasts seemingly surviving reprogramming but not becoming iPSCs (cluster 4). B. We used Seurat’s AddModuleScore command to average expression of 8-10 previously described markers of pluripotency as well as epithelial, mesenchymal, and fibroblast cell identity in each cell. On the UMAP, we recolored each cell for its score for each module. High expression is marked in red while low expression is marked in blue. C. Schematic demonstrating how to interpret different values for the mixing coefficient, previously described in (Goyal et al., 2021) (see Methods). Higher values of the mixing coefficient indicate a higher similarity in the expression profiles of the sets of barcoded cells analyzed. Representative UMAPs for different mixing coefficient values are shown. We calculated mixing coefficients for twins from the same clone on separate plates within the same reprogramming condition (in pink for iPSCs formed from OKSM alone, in blue for iPSCs formed from OKSM with LSD1 inhibition) and across reprogramming conditions (in gray). P-values comparing sample medians were calculated using the Wilcoxon rank sum test. D. On the UMAP, we recolored matched twins forming iPSCs with OKSM alone in pink and those forming iPSC with OKSM and LSD1 inhibition in light blue. For all clones with twins forming iPSCs in both reprogramming conditions, we measured the cluster containing the largest fraction of twins (i.e., dominant cluster) and calculated the fraction of all clones having that dominant cluster in each reprogramming condition. For each clone, we compared the dominant cluster in iPSCs formed with OKSM alone to the dominant cluster in iPSCs formed with OKSM and LSD1 inhibition, indicating a “switch” whenever the dominant cluster changed across reprogramming conditions. We measured the fraction of pairwise switches for each clone and marked them on the UMAP. The width of each arrow indicates the relative fraction of clones making that switch from OKSM alone to OKSM with LDS1 inhibition. E. For each matched pair twin across reprogramming conditions identified in Figure 3D, we plotted the average assigned pluripotency module scores for the twins reprogrammed in OKSM alone on the x-axis and for the twins reprogrammed in OKSM with LSD1 inhibition on the y-axis. Each dot represents an individual clone barcode. The p-value comparing paired means was calculated using a paired t-test. To measure differential expression of individual pluripotency markers, we calculated log2(fold change) values for each clone barcode and a subset of genes. We selected the pluripotency markers used in Figure 1 and also additional pluripotency markers associated with clusters 0, 1, and 3 found by running Seurat’s FindMarkers command. Metric shown is mean +/- 95% confidence interval for n = 26 clone barcodes. F. To identify clones forming iPSCs only when reprogrammed with OKSM and LSD1 inhibition (i.e., “LSD1i-dependent clones” in light blue) versus clones forming iPSCs in both reprogramming conditions (i.e., “LSD1i-independent clones” in dark blue), we performed barcode DNA-sequencing separately for each programming condition on the leftover iPSCs (see Figure S10C). We calculated the fraction of clone barcodes in each condition and plotted the barcode overlap as a Venn diagram with the percentage of all clone barcodes shown for each subset. On the UMAP, we plotted only cells for which we could confidently assign a single clone barcode and reprogrammed in OKSM with LSD1 inhibition. G. To visualize different axes of biological variability in our single-cell RNA-sequencing dataset, we aggregated expression profiles using Seurat’s AggregateExpression command for fibroblasts (cluster 4), indeterminate cells (cluster 5), iPSCs formed in OKSM alone (i.e., “OKSM alone iPSCs” in red), and iPSCs formed in OKSM with LSD1 inhibition from LSD1i-dependent clones (in light blue) versus from LSD1i-independent clones (in dark blue). We extracted loadings for each principal component and identified that the pluripotency markers used in Figure 5B had a significant contribution to principal component 1 in the leftward direction.

Having mapped the heterogeneous outcomes of reprogramming for normal induction with OKSM, we then set about measuring the differences in these outcomes that arise due to inhibition of LSD1 during induction. In order to track the outcomes of cells across these two conditions, after barcoding we let cells divide into multiple twins, which were then separated into four separate splits: two were subjected to standard OKSM induction and two were subjected to OKSM induction together with LSD1 inhibition. By comparing outcomes between twins across the same condition, one can determine the extent to which fates are predetermined, and comparing across different conditions reveals the differences in the composition and character of outcome states between those conditions. (Overall, we found that there were broadly minimal differences in the distribution of iPSCs, fibroblasts, and indeterminate cells between OKSM induction versus OKSM induction with LSD1 inhibition (Figure 5A).)

To measure the similarity of final molecular states both within and across clones, we used a previously formulated “mixing coefficient” metric. Briefly, the mixing coefficient measures how evenly interspersed distributions across different splits are in principal component space (Goyal et al., 2021); a mixing coefficient near 1 indicates high mixing or similarity while near 0 indicates low mixing or similarity (Figure 5C). Recent studies have shown that twins subjected to the same conditions can have either similar (Goyal et al., 2021; Richman et al., 2023) or distinct (Jiang et al., 2022) outcomes, suggesting a high degree of intrinsic or extrinsic fate specification, respectively. Here, within each reprogramming condition, we saw a high degree of mixing when comparing twins on separate plates (Figure 5C). However, we saw a similarly high degree of mixing between non-twins, indicating that within a set of reprogramming conditions clones are relatively mixed with no clear preference for distinct final states. To account for the relative low number of cells per clone, we performed a similar analysis with a more sensitive metric of similarity of single-cell molecular profiles (Jiang et al., 2022). While we found some amount of dissimilarity when comparing individual clones and equal numbers of randomly sampled cells, several clones continued to show no difference from random (Figure S9C).

We next wondered if twins across instead of within reprogramming conditions were transcriptionally similar. We found that matched twins in OKSM versus OKSM with LSD1 inhibition had a low degree of mixing, indicating distinct molecular profiles dictated by each respective reprogramming condition itself (Figure 5C). These results collectively suggest that what final state a cell ends up in after becoming an iPSC is dictated more by the extrinsic reprogramming conditions and less by pre-existing intrinsic differences before OKSM induction in our system.

Given that the same clones seemed to have distinct final states in iPSCs formed from OKSM alone versus OKSM with LSD1 inhibition, we wondered what was different about those final states. We compared matched twins from the same clone across each condition. A significant fraction of clones primarily in cluster 0 in OKSM induction alone “switched” to primarily be in cluster 3 (60%) or cluster 1 (25%) in OKSM induction with LSD1 inhibition (Figure 5D). To broadly assess the pluripotency status of these cells, we assigned a pluripotency module score to each cell based on averaged expression of 10 commonly used pluripotency markers and found no significant difference between matched twins across conditions (Figure 5E).

Looking at specific pluripotency markers individually, however, iPSCs demonstrated heterogeneity within and across conditions, as previously seen in iPSC culture (Masaki et al., 2007; Narsinh et al., 2011). We saw minimal differences in expression of the core pluripotency factors *NANOG* and *OCT4*. Among markers selected for the pluripotency module, DNA methyltransferase *DNMT3B* and RNA-binding protein *LIN28A*, both with known roles in pluripotency maintenance, were modestly elevated in twins in OKSM induction alone.

Meanwhile, *PODXL*, a surface marker used for isolating “functional” iPSCs, was modestly elevated in twins in OKSM induction with LSD1 inhibition (Cai et al., 2006). We further identified a number of genes less commonly associated with pluripotency but strongly correlated with our gene expression clusters; twins forming iPSCs in OKSM induction with LSD1 inhibition had significantly higher levels of *NLRP7*, known to block BMP4-mediated downregulation of pluripotency factors and differentiation, and lower levels of *TERF1*, known to promote telomere elongation and pluripotency factor expression, when compared to twins forming iPSCs in OKSM induction alone (Alici-Garipcan et al., 2020; Q. Liu et al., 2018). These results indicate that LSD1 inhibition may push reprogramming cells towards certain final iPSC states marked by differential expression of several pluripotency markers not reported in previous bulk RNA-sequencing analyses (Cacchiarelli et al., 2015).

LSD1 inhibition (LSD1i) during iPSC reprogramming increased the number of primed clones and consequently the number of iPSC colonies formed by “reclassifying” previously nonprimed clones as primed (Figure 4). We wondered whether clones reclassified from nonprimed to primed with LSD1 inhibition (i.e., LSD1i-dependent) could be distinguished from clones classified as primed without or with LSD1 inhibition (i.e., LSD1i-independent) after becoming iPSCs. To identify LSD1i-independent and -dependent clone barcodes while avoiding significant subsampling, we performed barcode DNA-sequencing and measured barcode overlap as in Figure 4B on the leftover reprogrammed iPSCs in each condition after removing a small fraction for single-cell RNA-sequencing (Figure 5F). The LSD1i-dependent clones were, however, essentially indistinguishable from the LSD1i-independent clones when comparing gene expression profiles. When we visualized aggregates of each category in principal component space, we found iPSCs formed via reprogramming with OKSM and LSD1 inhibition derived from LSD1i-independent and from LSD1i-dependent clones to be virtually overlapping (Figure 5G). The positioning of different iPSC subpopulations in principal component space further reiterated that LSD1 inhibition during reprogramming creates iPSCs that are subtly distinct from those formed by OKSM alone.

In sum, while pre-existing molecular differences strongly influence whether a cell becomes an iPSC or not, such differences seemingly have much less influence on the final transcriptional state of the resultant iPSCs, reflecting either a lack of memory of those initial differences or intrinsic homogeneity of the iPSC fate itself.

## Discussion

While much work has gone into identifying and characterizing the molecular characteristics and mechanisms of cells *en route* to becoming iPSCs following induction of OKSM, the demonstration that “twins” share the same reprogramming outcome (Pour et al., 2015; Shakiba et al., 2019; Yunusova et al., 2017) suggested the existence of intrinsic differences in otherwise homogeneous-seeming cells that drive reprogramming outcomes. Rewind, one of a number of tools that enable one to retrospectively connect fates to initial primed states, allowed us to characterize these primed states directly (Emert et al., 2021; Tian et al., 2018; Umkehrer et al., 2021; Weinreb et al., 2020). Here, we revealed that primed fibroblasts are marked by naturally-arising fast cycling speed and lack of fibroblast activation, both of which likely represent a single underlying axis of biological variability and both of which directly affect iPSC reprogramming efficiency.

Although Rewind was able to reveal new information about cells primed for reprogramming, it is worth noting that there are likely additional important features to be discovered. For instance, while cycling speed and fibroblast activation emerged as the strongest measureable priming features in our analyses, in logistic regression models each variable alone only explained 0.5%-1% of variation in priming and when combined as a principal component only explained 9.3% of variation in gene expression, leaving open the question of what explains the remaining variability—indeed, there were already hints of another principal component that may additionally mark the primed state (Figure 3D). One source of variability is undoubtedly technical: given that the primed cells are a very rare subset, we often had only 10s-100s of primed cells for our analyses, leaving us with a high degree of noisiness in our measurements. Other sources of variability that could affect the Rewind methodology could be a loss of memory between the cell divisions required for the technique to work.

A major question raised by our work is how to define the primed state. It is tempting to use a purely cell-intrinsic definition of priming consisting of a discrete set of molecular markers. However, as our results using the LSD1 inhibitor during reprogramming show, cells that are not primed for reprogramming in one condition can be primed for reprogramming in another. Hence, any definition must incorporate both the molecular state of the cell as well as the stimulus applied. We have observed similar reclassifications in other systems when different perturbation conditions are applied (Emert et al., 2021; Goyal et al., 2021; Torre et al., 2021), suggesting that such a definition for priming may be required in other contexts as well.

The fact that cells can be primed in one condition and not primed in another also eliminates the possibility that there is a single, distinct primed molecular state. Rather, there must be multiple primed states, given that different cells can have different outcomes in different conditions. Our prior work has demonstrated the existence of multiple primed states in the context of cancer therapy resistance (Dardani et al., 2022; Emert et al., 2021; Goyal et al., 2021; Torre et al., 2021). Are these primed states organized along a single axis of variability, or multiple? In the case of cancer therapy resistance, we found evidence for multiple axes of variability (Dardani et al., 2022; Emert et al., 2021). Another question is whether primed cell states form a continuum, or if they consist of discrete metastable states that cells can fluctuate between (Chang et al., 2008; Mojtahedi et al., 2016; Schuh et al., 2020). It is difficult to answer this question with current approaches. New conceptual approaches (experimental and theoretical) for the identification of metastable states will be required to provide answers.

Another question raised by our work is whether priming constitutes a “head start” towards a target cell type or rather a deviation to a generically primed state in which the cell is capable of a number of fate transitions. Our data would argue against priming being a head start in this instance, given that there is virtually no difference in expression of any pluripotency markers between primed and nonprimed cells (Wakao et al., 2011). Whether the primed cells we found are primed for other cell fate transitions is an important future direction.

A reasonable alternative to priming constituting a “head start” could be that primed cells exhibit some degree of de-differentiation. We observed that fibroblast markers were unchanged in primed cells (versus nonprimed cells), but primed cells did express comparatively lower levels of markers of fibroblast activation and myofibroblast differentiation. The hiF-T cell line is a secondary fibroblast line (meaning that it was formed from reprogramming fibroblasts into iPSCs and then differentiating a clone back into fibroblast-like cells), so it is unclear how to interpret the expression of these markers in this line, which may arise from the culturing conditions used (Baranyi et al., 2019; Baum & Duffy, 2011; Doolin et al., 2021; Hinz et al., 2012; López-Antona et al., 2022; Masur et al., 1996; Pakshir et al., 2020). The literature is also mixed on the effects of fibroblast activation on iPSC reprogramming, with some studies reporting that activated fibroblasts and myofibroblasts taken from fibrotic or damaged tissues are known to be more resistant to iPSC reprogramming (Song et al., 2016; Tanaka et al., 2020) while others reporting that mouse fibroblasts with an enhanced propensity to form myofibroblasts in uninduced populations reprogram into iPSCs with higher efficiency (Koumas et al., 2003; Nemajerova et al., 2012; Sanders et al., 2007). Together, we think it is difficult to say whether primed cells are truly de-differentiated or just in some other fibroblast state.

Cell fate is typically thought to be determined by some combination of cell intrinsic and cell extrinsic factors. Intrinsic factors can be defined as those within the cell that, if the cell were placed in a different environment, would still dictate the same outcome, whereas extrinsic factors are those environmental factors that would dictate the outcome regardless of the internal state of the cell. An important concept to introduce here is that of memory, namely how long the intrinsic determination persists over time. Schematically, the following classification may hold: (1) Cell fate is determined by extrinsic factors (Biddy et al., 2018; Jiang et al., 2022); (2) Cell fate is determined by very short-lived intrinsic factors, thereby seeming “stochastic” (Luria & Delbrück, 1943; Symmons & Raj, 2016); (3) Cell fate is determined by intermediate-lifetime intrinsic factors, in which case close progeny may adopt the same fate as the ancestor (Emert et al., 2021; Goyal et al., 2021; Shaffer et al., 2020; Weinreb et al., 2020); (4) Cell fate is determined by very long-lived intrinsic factors (e.g., mutations), in which case all progeny will adopt the same fate as the ancestor. With the advent of barcoding techniques, it has become possible to rigorously distinguish between these possibilities. In the case of reprogramming, much ink has been spilled on the distinction between extreme cases 2 and 4, i.e., either completely stochastic induction versus completely deterministic induction, with both cases seemingly supported and contradicted by experimental evidence. We propose that case 3, an intermediate level of memory, may be a way to reconcile the various data. In that scenario, cells have a limited memory of the primed state, so once a cell enters the primed state, its progeny will eventually “forget”. On a long time scale, such as expanding individual cells into large clonal populations (J. Hanna et al., 2009), this loss of cellular memory would make priming for reprogramming seem stochastic, but on shorter timescales, such as the few divisions used in ours and others’ barcoding experiments, priming will be inherited and seem deterministic. Such intermediate memory timescales have been found in many other systems (Emert et al., 2021; Mold et al., 2022; Shaffer et al., 2017, 2020). The mechanisms underlying these intermediate forms of memory remain to be revealed.

Given that there is a molecular distinction between cells that were primed to reprogram versus those that were not, it was conceivable that smaller differences even between individual primed cells could propagate to differences between the resultant iPSCs. With the advent of barcoding technologies, this form of clonal memory has revealed itself in a number of systems (Goyal et al., 2021; Mold et al., 2022; Richman et al., 2023; Tian et al., 2018; Umkehrer et al., 2021). In the reprogramming system we analyzed here, we found little evidence for clonal memory in the final state, with iPSCs seeming largely the same even when we knew there were differences in the initial state (e.g., LSD1i-dependent and -independent primed cells formed indistinguishable iPSCs). It could be that measurement techniques based solely on RNA cannot reveal the differences driven by clonal memory. Alternatively, it could be that iPSCs represent an attractor state that erases the molecular history of the cell.

## Methods

### Cell lines and culture conditions

Unless otherwise noted, all cell culture incubations were performed at 37°C, 5% CO_2_. We tested intermittently for mycoplasma contamination. We cultured hiF-T cells as previously described prior to hiF-T reprogramming experiments (Cacchiarelli et al., 2015). Briefly, we expanded hiF-T cells in growth medium on TC plastic dishes coated with Attachment Factor (Fisher #S006100), and split cells 1:3 upon reaching 60%-70% confluency. hiF-T growth medium (GM) is DMEM/F-12 with Glutamax (Life Tech #10585018) + 10% ES-FBS (Life Tech #16141079) + 1X 2-mercaptoethanol (Life Tech #21985023) + 1X NEAA (Invitrogen #11140050) + P/S + 0.5ug/mL puromycin + 16 ng/mL rhFGF-basic (Promega #G5071). When passaging hiF-T cells, we performed dissociation with accutase (Sigma #A6964-100ML), and followed the manufacturer’s instructions.

### Reprogramming to pluripotency

We performed hiF-T reprogramming experiments as previously described (Cacchiarelli et al., 2015; Mellis et al., 2021). Briefly, we expanded hiF-T cells in hiF-T GM without puromycin for one week. On day -1, we seeded CF-1 irradiated MEFs (Fisher #A34181) on 24-well plates (Corning #353047) coated with Attachment Factor (Fisher #S006100). On day 0, we seeded varying amounts (1-3 * 10^5) hiF-T cells per 24-well plate. On day 1, we began Yamanaka factor induction by switching media to hiF-T GM with 2 ug/mL doxycycline and without puromycin. On day 3, we switched media to KSR medium (KSRM): DMEM/F-12 with Glutamax (Life Tech #10585018) + 20% Knockout Serum Replacement (Life Tech #10828010) + 1X 2-mercaptoethanol (Life Tech #21985023) + 1X NEAA (Invitrogen #11140050) + P/S + 8 ng/mL rhFGF-basic (Promega #G5071) + 2 ug/mL doxycycline. We changed KSRM daily, and analyzed cells on day 21. We performed ≥2 biological replicates (i.e. different vials of hiF-T cells expanded and reprogrammed on different days with different batches of media) unless otherwise specified for all experiments. For reprogramming experiments with perturbations, we used the LSD1 inhibitor RN-1 (Millipore #489479) at a final concentration of 1uM and the DOT1L inhibitor pinometostat (Selleck Chemicals #S7062) at a final concentration of 4uM.

### Alkaline phosphatase staining with colorimetry

We used the Vector Red Substrate kit (Vector Labs #SK-5100) to stain hiF-T-iPSC colonies after fixation on day 21 of reprogramming experiments. We fixed wells in 24-well format using 3.7% formaldehyde for 3 min, and followed the manufacturer’s instructions.

### Embryoid body formation from hiF-T-iPSC colonies

After forming hiF-T-iPSC colonies following 21 days of OKSM induction, we evaluated differentiation of hiF-T-iPSCs into embryoid bodies. We dissociated hiF-T-iPSCs from MEF feeder cells with accutase (Sigma #A6964-100ML) for 2-10 min at 37°C before adding embryoid body (EB) media: DMEM/F-12 with Glutamax (Life Tech #10585018) + 20% ES-FBS (Life Tech #16141079) + 1X 2-mercaptoethanol (Life Tech #21985023) + 1X NEAA (Invitrogen #11140050) + P/S. We dislodged colonies and mechanically broke up colonies by pipetting up and down to form a cell suspension. We performed two washes with centrifugation at 1200 rpm for 2 min and resuspension in 5 mL of EB media. Because MEF feeders and hiF-T-iPSCs both dissociated with accutase, we briefly placed the cell suspension on 10 cm dishes coated with Attachment Factor (Fisher #S006100) for 30 min at 37°C to enable preferential adhering of the MEF feeders but not hiF-T-iPSCs. The hiF-T-iPSCs largely remained in suspension, which we collected. We plated 1 mL of final cell suspension into each well of an ultra-low attachment surface 6-well plate (Corning #3471) and allowed the hiF-T-iPSCs to aggregate and differentiate into embryoid bodies over 14 days. To help with cell survival, we incubated cells in 10uM of ROCK inhibitor Y26632 (Calbiochem #688001) for the first 2 days. We replaced the media every 2 days by collecting aggregates into a 15 mL conical, incubating for 15-30 min at 37°C to allow the aggregates to settle at the bottom, and carefully replacing the media. On day 14, we collected barcoded embryoid bodies for DNA-sequencing of clone barcodes and plated a small amount on normal 6-well plates to enable adhering and differentiation of embryoid bodies into different cell types, which we confirmed by light microscopy imaging.

### Clone barcode library lentivirus generation

Barcode libraries were constructed as previously described (Emert et al., 2021; Goyal et al., 2021; Jiang et al., 2022; Torre et al., 2021). Full protocol available at https://www.protocols.io/view/barcode-plasmid-library-cloning-4hggt3w. Briefly, we modified the LRG2.1T plasmid (kindly provided by Junwei Shi) by removing the U6 promoter and single guide RNA scaffold and inserting a spacer sequence flanked by EcoRV restriction sites just after the stop codon of GFP. We digested this vector backbone with EcoRV (NEB #R3195S) and gel purified the resulting linearized vector. We ordered PAGE-purified ultramer oligonucleotides (IDT) containing 30 nucleotides homologous to the vector insertion site surrounding 100 nucleotides with a repeating “WSN” pattern (W = A or T, S = G or C, N = any) and used Gibson assembly followed by column purification to combine the linearized vector and barcode oligo insert. We performed electroporations of the column-purified plasmid into Endura electrocompetent *E. coli* cells (Lucigen #60242-1) using a Gene Pulser Xcell (Bio-Rad #1652662), allowing for recovery before plating serial dilutions and seeding cultures (200 mL each) for maxipreparation. We incubated these cultures on a shaker at 225 rpm and 32 °C for 12–14 h, after which we pelleted cultures by centrifugation and used the EndoFree Plasmid Maxi Kit (Qiagen #12362) to isolate plasmid according to the manufacturer’s protocol, sometimes freezing pellets at −20°C for several days before isolating plasmid. Barcode insertion was verified by polymerase chain reaction (PCR) from colonies from plated serial dilutions. We pooled the plasmids from the separate cultures in equal amounts by weight before packaging into lentivirus. We estimated our library complexity as described elsewhere (Goyal et al., 2021). Briefly, we sequenced three independent transductions in WM989 A6-G3 melanoma cells and took note of the total and pairwise overlapping extracted barcodes. Using the mark-recapture analysis formula, we estimate our barcode diversity from these three transductions to be between 48.9 and 63.3 million barcodes.

### Lentivirus packaging of clone barcode library

We adapted previously described protocols to package lentivirus (Emert et al., 2021; Goyal et al., 2021; Jiang et al., 2022; Torre et al., 2021). We first grew HEK293FT to near confluency (80-95%) in 10 cm plates in DMEM + 10% FBS + P/S. On day -1, we changed the media in HEK293FT cells to DMEM + 10% FBS without antibiotics. For each 10 cm plate, we added 80 µL of polyethylenimine (Polysciences #23966) to 500 µL of Opti-MEM (Fisher #31985062), separately combining 5 µg of VSVG and 7.5 µg of pPAX2 and 7.35 µg of the barcode plasmid library in 500 µL of Opti-MEM. We incubated both solutions separately at room temperature for 5 min. Then, we mixed both solutions together by vortexing and incubated the combined plasmid-polyethylenimine solution at room temperature for 15 min. We added 1.09 mL of the combined plasmid-polyethylenimine solution dropwise to each 10 cm dish. After 6-7 hours, we aspirated the media from the cells, washed with 1X DPBS, and added fresh hiF-T GM. The next morning, we aspirated the media, and added fresh hiF-T GM. Approximately 9-11 hours later, we transferred the virus-laden media to an empty, sterile 50 mL conical tube and stored it at 4°C, and added fresh hiF-T GM to each plate. We continued to collect the virus-laden media every 9-11 hours for the next 30 hours in the same 50 conical mL tube, and stored the collected media at 4°C. Upon final collection, we filtered the virus-laden media through a 0.45µm PES filter (Millipore-Sigma #SE1M003M00) and stored 1.5 mL aliquots in cryovials at -80ºC.

### Transduction with lentiviral clone barcode library

To transduce hiF-T cells, we freshly thawed virus-laden media on ice, added it to dissociated cells with 4ug/mL of polybrene (Millipore-Sigma #TR-1003-G), and plated 50,000 cells/well in a 6-well plate coated with Attachment Factor (Fisher #S006100). The volume of virus-laden media used was decided by measuring the multiplicity of infection (MOI) with different viral titers. For single-cell RNA-sequencing experiments, we aimed for a low MOI with 10%-25% GFP-positive cells to minimize the fraction of cells with multiple unique barcodes. We found it relatively computationally challenging to differentiate multiple-barcoded cells from doublets introduced by gel beads-in-emulsions. This was not an issue for bulk DNA clone barcode overlap experiments for which we aimed for a high MOI with 60-70% GFP-positive cells. We used 35 uL/well of virus-laden media for low MOI and 80 uL/well for high MOI. After plating hiF-T cells with virus, we performed a 30 min incubation at room temperature before centrifuging the 6-well plate at 930g for 30 min at room temperature. After 24 hours, we passaged the cells from 2 wells onto 10 cm plates or from 6 wells onto 15cm plates. The barcoded cells (GFP-positive) were sorted for all single-cell RNA-sequencing experiments but not for bulk DNA clone barcode overlap experiments.

### DNA-sequencing of clone barcodes from genomic DNA

We prepared clone barcode sequencing libraries from genomic DNA (gDNA) as previously described (Emert et al., 2021; Goyal et al., 2021; Jiang et al., 2022). Briefly, we isolated gDNA from barcoded cells using the QIAmp DNA Mini Kit (Qiagen #51304) per the manufacturer’s protocol. Extracted gDNA was stored as a pellet in -20ºC for days to weeks before the next step. We then performed targeted amplification of the barcode using custom primers containing Illumina adaptor sequences, unique sample indices, variable-length staggered bases, and an “UMI” consisting of 6 random nucleotides (NHNNNN). As reported in (Emert et al., 2021), the “UMI” does not uniquely tag barcode DNA molecules, but nevertheless appeared to increase reproducibility and normalize raw read counts. We determined the number of amplification cycles (N) by initially performing a separate quantitative PCR (qPCR) and selecting the number of cycles needed to achieve one-third of the maximum fluorescence intensity for serial dilutions of genomic DNA. The thermocycler (Veriti #4375786) was set to the following settings: 98°C for 30 sec, followed by N cycles of 98°C for 10 sec and then 65°C for 40 sec and, finally, 65 °C for 5 min. Upon completion of the PCR reaction, we immediately performed a 0.7X bead purification (Beckman Coulter SPRISelect #B23319), followed by final elution in nuclease-free water. Purified libraries were quantified with a High Sensitivity dsDNA kit (Thermo Fisher #Q33230) on a Qubit Fluorometer (Thermo Fisher #Q33238), pooled, and sequenced on a NextSeq 500 machine (Illumina) using 150 cycles for read 1 and 8 reads for each index (i5 and i7). The primers used are described in (Emert et al., 2021; Goyal et al., 2021).

### Analyses of sequenced clone barcodes from genomic DNA

The barcode libraries from genomic DNA-sequencing data were analyzed as previously described (Emert et al., 2021), with the custom barcode analysis pipeline available at https://github.com/arjunrajlaboratory/timemachine. Briefly, this pipeline searches for barcode sequences that satisfy a minimum phred score and a minimum length. We also use STARCODE (Zorita et al., 2015), available at https://github.com/gui11aume/starcode, to merge sequences with Levenshtein distance ≤ 8 and add the counts across collapsed (merged) barcode sequences. To normalize reads for differently indexed samples and correct for sequencing depth, we calculated reads per million (RPM) for each barcode. This normalization was insufficient when comparing iPSCs reprogrammed in OKSM alone versus in OKSM with perturbations to increase overall efficiency (i.e., inhibition of LSD1 or DOT1L). When reprogramming in OKSM with perturbations a higher fraction of clones form iPSCs, so each individual clone has less representation and hence reads even when correcting for sequencing depth. For our early clone barcode sequencing experiments (Figures 1C, 1E, 4B-C, S2A-D, S8), we performed an additional correction by assuming extremely large iPSC colonies appearing in both conditions had similar representation in each. We determined the average fold change for the largest 5-10 shared iPSC colonies between each condition and multiplied all RPM values for each barcode in the OKSM with perturbation conditions by this fold change value. Additionally, we made use of two subclones (D8 and F8) of WM989 A6-G3 generated in (Goyal et al., 2021) as spike-in standards to convert sequencing counts into relative cell numbers (Figures 3A, 4C, 5A, 5F, S8, S10C). We spiked in a known number of cells for each barcoded subclone to each cell pellet before gDNA extraction and sequencing. Then, we used linear regression (on (0,0), (count_F8, cells_F8), (count_D8, cells_D8) to get the conversion factor from read counts of all barcodes to cell numbers. We used a minimum cell count (determined either by large colony or spike-in normalization) and log2-fold change between pairs of conditions to annotate clones as condition-dependent or condition-independent. We obtained similar results when using each normalization method (see Figure S8).

### Simulation for clone barcode overlap

We adapted a described previously computational model that simulates all steps of our experiments designed to compare barcode overlap in resistant colonies (Yunusova et al., 2017). The model simulates cell seeding and infection. Each cell is represented as an independent object. The number of barcoded cells was calculated as number of barcoded cells = number of seeded cells x (1 - e^-MOI), where the MOI was estimated for our barcode lentivirus. Barcodes were represented by integer numbers from among 20 million variants of unique barcodes as a conservative estimate from our lentiviral library diversity (Emert et al., 2021; Goyal et al., 2021). The subset of barcoded cells was assigned barcodes randomly with replacement from this library. The model simulates expanding cells prior to induction of OKSM. Each cell, regardless of barcode status, undergoes a cell division procedure with 2-4 rounds depending on the experimental condition. In each round, a given cell will give rise to a number of progeny sharing the same barcode based on an estimated distribution of cell division in hiF-T cells. The model plates cells onto separate dishes/splits (with number of dishes/splits dependent on the experiment) by randomly assigning each cell an integer. The model simulates the formation of resistant colonies assuming a purely stochastic model of iPSC reprogramming. A defined fraction of cells on each plate form iPSC colonies based on a reprogramming efficiency that was calculated as reprogramming efficiency = number of barcoded iPSC colonies after reprogramming / number of seeded barcoded cells based on experimental observations. Additionally, each cell forming an iPSC colony is subject to probabilistic material loss at different stages of the *in silico* experiment, including cell culture (5%) and genomic DNA extraction (5%). The output of the model was the number of barcodes shared between different plates or barcode overlap. This was not corrected for cells having more than one lentiviral barcode due to multiple integrations for a given MOI. We performed 1000 independent simulations to obtain a distribution of barcode overlap values to determine the probability of obtaining our observed barcode overlap from our experiments by random chance. This model was written and executed in R.

### Flow sorting of barcoded cells

We used accutase (Sigma #A6964-100ML) to detach the barcoded cells from the plate and subsequently neutralized the accutase with the corresponding media depending on the cell type (hiF-T GM for fibroblasts, KSRM for iPSCs). We then pelleted the cells, performed a wash with 1X DPBS (Invitrogen #14190-136), and resuspended again in 1X DPBS. Cells were sorted on a BD FACSJazz machine (BD Biosciences) or MoFlo Astrios machine (Beckman Coulter), gated for positive GFP signal and singlets. Sorted cells were then centrifuged to remove the supernatant media containing PBS, and either replated with the appropriate cell culture media or prepared for DNA-sequencing or RNA-sequencing.

### Single-cell RNA-sequencing

We used the 10x Genomics single-cell RNA-seq kit v3 to sequence barcoded cells. We resuspended the cells (aiming for up to 10,000 cells for recovery/ sample) in PBS and followed the protocol for the Chromium Next GEM Single Cell 3ʹ Reagent Kits v3.1 as per manufacturer directions (10x Genomics). Briefly, we generated gel beads-in-emulsion (GEMs) using the 10x Chromium system, and subsequently extracted and amplified barcoded cDNA as per post-GEM RT-cleanup instructions. We then used a fraction of this amplified cDNA (25%) and proceeded with fragmentation, end-repair, poly A-tailing, adapter ligation, and 10x sample indexing per the manufacturer’s protocol. We quantified libraries using the High Sensitivity dsDNA kit (Thermo Fisher #Q32854) on Qubit 2.0 Fluorometer (Thermo Fisher #Q32866) and performed Bioanalyzer 2100 (Agilent #G2939BA) analysis prior to sequencing on a NextSeq 500 machine (Illumina) using 28 cycles for read 1, 55 cycles for read 2, and 8 cycles for i7 index.

### Analyses of expression data from single-cell RNA-sequencing

We adapted the cellranger v3.0.2 by 10x Genomics into our custom pipeline (https://github.com/arjunrajlaboratory/10xCellranger) to map and align the reads from the NextSeq sequencing run. Briefly, we downloaded the BCL counts and used cellranger mkfastq to demultiplex raw base call files into library-specific FASTQ files. We aligned the FASTQ files to the hg38 human reference genome and extracted gene expression count matrices using cellranger count, while also filtering and correcting cell identifiers and unique molecular identifiers (UMI) with default settings. We then performed the downstream single-cell expression analysis in Seurat v3. Within each experimental sample, we removed genes that were present in less than three cells, as well as cells with less than or equal to 200 genes. We also filtered for mitochondrial gene fraction which was dependent on the cell type. For non-identically treated samples, we integrated them using scanorama (Hie et al., 2019), which may work better to integrate non-similar datasets and avoid over-clustering. For samples that were exposed to identical treatment, we normalized using SCTransform (Hafemeister and Satija, 2019) and the samples according to the Satija lab’s integration workflow (https://satijalab.org/seurat/articles/integration_introduction.html). For each experiment, we used these integrated datasets to generate data dimensionality reductions by principal component analysis (PCA) and Uniform Manifold Approximation and Projection (UMAP), using 50 principal components for UMAP generation. For a majority of analyses, we worked with the principal component space and normalized expression counts. To determine what resolution for clustering to use for each single-cell RNA-sequencing dataset, we used the clustree package in R (Zappia & Oshlack, 2018). Briefly, we visualized the relationships between clusters at multiple resolutions (0 - 1 in steps of 0.1) and chose the highest resolution before individual cluster nodes began to have multiple incoming edges, which indicates overclustering. We used a resolution of 0.45 for our dataset in Figure 1, a resolution of 0.3 for our dataset in Figure 2, and a resolution of 0.3 for our dataset in Figure 5. In several of our single-cell RNA-sequencing datasets, we noticed several outlier clusters with lower numbers of counts and numbers of features usually combined with higher percentages of mitochondrial or ribosomal counts in comparison to the majority of clusters. We decided to remove said clusters from our datasets before performing further analyses. These outlier clusters corresponded to clusters 4, 9, and 10 for our dataset in Figure 1 and to clusters 6, 7, and 8 for our dataset in Figure 5.

### Clone barcode recovery from single-cell RNA-sequencing libraries

As the clone barcodes are transcribed, we extracted the barcode information from the amplified cDNA from the 10x Genomics V3 chemistry protocol. We ran a PCR side reaction with one primer that targets the 3’ UTR of GFP and one primer that targets a region introduced by the amplification step within the V3 chemistry protocol (“Read 1”). The two primers amplify both the 10x cell-identifying sequence as well as the 100 base pair barcode that we introduced lentivirally. The number of cycles, usually between 12-15, was decided by calculating the cycle threshold value from a qPCR reaction using the NEBNext Q5 Hot Start HiFi PCR Master Mix (NEB #M0543L) for the specified cDNA concentration. The thermocycler (Veriti #4375305) was set to the following settings: 98°C for 30 sec, followed by N cycles of 98°C for 10 sec and then 65°C for 2 min and, finally, 65°C for 5 min. Upon completion of the PCR reaction, we immediately performed a 0.7X bead purification (Beckman Coulter SPRISelect #B23319) followed by final elution in nuclease-free water. Purified libraries were quantified with High Sensitivity dsDNA kit (Thermo Fisher #Q33230) on Qubit Fluorometer (Thermo Fisher #Q33238), pooled, and sequenced on a NextSeq 500 machine (Illumina). We sequenced 26 cycles on Read 1 which gives 10x cell-identifying sequence and UMI, 124 cycles for read 2 which gives the barcode sequence, and 8 cycles for index i7 to demultiplex pooled samples. The primers used are described in (Goyal et al., 2021).

### Analyses of clone barcode data from single-cell RNA-sequencing

The barcodes from the side reaction of single-cell cDNA libraries were recovered by developing custom shell, R, and Python scripts, which are all available at this link: https://github.com/arjunrajlaboratory/10XBarcodeMatching. Briefly, we scan through each read searching for sequences complementary to the side reaction library preparation primers, filtering out reads that lack the GFP barcode sequence, have too many repeated nucleotides, or do not meet a phred score cutoff. Since small differences in otherwise identical barcodes can be introduced due to sequencing and/or PCR errors, we merged highly similar barcode sequences using STARCODE software (Zorita et al., 2015), available at https://github.com/gui11aume/starcode. For varying lengths of barcodes (30, 40 or 50) depending on the initial distribution of Levenshtein distance of non-merged barcodes, we merged sequences with Levenshtein distance ≤ 8, summed the counts, and kept only the most abundant barcode sequence. For downstream analysis, we filtered out all barcodes that were associated below a conservative minimum cutoff (dependent on sequencing depth) of unique molecular identifiers (UMI). For cases where one 10x cell-identifying sequence was associated with more than one unique barcode, we calculated the fraction of UMIs per barcode for each cell and filtered out all barcodes that did not make up at least 40% of UMIs. Finally, we filtered out all cells that were still associated with more than one unique barcode after these filtering steps. This could either result from multiplets introduced within gel beads-in-emulsions or because of the same cell receiving multiple barcodes during lentiviral transduction. We were able to confidently recover barcodes associated with 30-40% of single cells, which were then used in the downstream clone-resolved analysis.

### Single-molecule RNA FISH on cells in plates

We performed single-molecule RNA FISH as previously described (Raj et al., 2008). For the genes used here, we designed complementary oligonucleotide probe sets using custom probe design software in MATLAB and ordered them with a primary amine group on the 3′ end from Biosearch Technologies (seeSupplementary Table 1 for probe sequences). We then pooled each gene’s complementary oligos and coupled the set to Cy3 (GE Healthcare), Alexa Fluor 594 (Life Technologies) or Atto 647N (ATTO-TEC) N-hydroxysuccinimide ester dyes. The cells were fixed as follows: we aspirated media from the plates containing cells, washed the cells once with 1X DPBS, and then incubated the cells in the fixation buffer (3.7% formaldehyde in 1X DPBS) for 10 min at room temperature. We then aspirated the fixation buffer, washed samples twice with 1X DPBS, and added 70% ethanol before storing samples at 4ºC. For hybridization of RNA FISH probes, we rinsed samples with wash buffer (10% formamide in 2X SSC) before adding hybridization buffer (10% formamide and 10% dextran sulfate in 2X SSC) with standard concentrations of RNA FISH probes and incubating samples overnight with coverslips, in humidified containers at 37ºC. The next morning, we performed two 30 min washes at 37ºC with the wash buffer, after which we added 2X SSC with 50 ng/mL of DAPI. We mounted the samples for imaging in 2X SSC. To strip RNA FISH probes to re-hybridize and re-image for additional genes, we incubated samples in stripping buffer (60% formamide in 2X SSC) for 20 min on a hot plate at 37ºC, washed samples 3 × 15 min with 1X PBS on a hot plate at 37ºC, then returned samples to 2X SSC. After stripping RNA FISH probes, we re-imaged all previous positions and excluded dyes with residual signal from subsequent hybridization.

### *Detecting clone barcodes in carbon copies* in situ

We adapted the Hybridization Chain Reaction (HCR V3.0) (Choi et al., 2018) for barcode RNA FISH as follows. We used 1.2 pmol each of up to 240 barcode RNA FISH probes per 0.3 mL hybridization buffer. Our primary hybridization buffer consisted of 30% formamide, 10% dextran sulfate, 9 mM citric acid pH 6.0, 50 μg/mL heparin, 1X Denhardt’s solution (Life Technologies #750018) and 0.1% Tween-20 in 5X SSC. For primary hybridization, we used 100 μL hybridization buffer per well of a 2 well plate, covered the well with a glass coverslip, and incubated the samples in humidified containers at 37ºC for 8 hours. Following the primary probe hybridization, we washed samples 4 × 5 min at 37ºC with washing buffer containing 30% formamide, 9 mM citric acid pH 6.0, 50 μg/mL heparin, and 0.1% tween-20 in 5X SSC. We then washed the samples at room temperature 2 × 5 min with 5X SSCT (5X SSC + 0.1% Tween-20) and incubated the samples at room temperature for 30 min in amplification buffer containing 10% dextran sulfate and 0.1% Tween-20 in 5X SSC. During this incubation, we snap-cooled individual HCR hairpins (Molecular Instruments) conjugated to Alexa Fluor 647 (Alexa647) by heating to 95ºC for 90 sec then immediately transferring to room temperature to cool for 30 min while concealed from light. After these 30 min, we resuspended and pooled the hairpin in amplification buffer to a final concentration of 6-15 nM each. We added the hairpin solution to samples along with a coverslip and incubated samples at room temperature overnight (12–16 hours) concealed from light. The following morning, we washed samples 5 × 5 min with 5X SSCT containing 50 ng/mL DAPI, added SlowFade antifade solution (Life Technologies #S36940) and a coverslip before proceeding with imaging. To remove fluorescent signal for subsequent rounds of RNA FISH or immunofluorescence, we photobleached samples on the microscope or stripped HCR hairpins as described above for RNA FISH probes.

### Imaging RNA FISH and colorimetric dye signal

To image single-molecule RNA FISH, nuclei, and colorimetric dye signal, we used a Nikon TI-E inverted fluorescence microscope equipped with a SOLA SE U-nIR light engine (Lumencor), a Hamamatsu ORCA-Flash 4.0 V3 sCMOS camera, and 4X Plan-Fluor DL 4XF (Nikon #MRH20041/MRH20045), 10X Plan-Fluor 10X/0.30 (Nikon #MRH10101) and 60X Plan-Apo λ (#MRD01605) objectives. We used the following filter sets to acquire signal from different fluorescence channels: 31000v2 (Chroma) for DAPI, 41028 (Chroma) for Atto 488, SP102v1 (Chroma) for Cy3, 17 SP104v2 (Chroma) for Atto 647N, and a custom filter set for Alexa Fluor 594. We tuned the exposure times depending on the dyes used (Cy3, Atto 647N, Alexa Fluor 594, and DAPI). For large scans, we used a Nikon Perfect Focus system to maintain focus across the imaging area. For imaging RNA FISH signal at high magnification (≥60X), we acquired z-stacks of multiple Z-planes and used the maximum intensity projection to visualize the signal.

### Computationally processing fluorescence microscopy images

For colony counting, Nikon-generated ND2 files were stitched and converted from ND2 format to TIFF format within the Nikon NIS-Elements software. The number of colonies within each sample was counted using custom MATLAB code, available at https://github.com/arjunrajlaboratory/blobCounter. To identify cells containing primed clone barcodes by RNA FISH, we used custom MATLAB scripts to stitch, contrast and compress scan images (scripts available at https://github.com/arjunrajlaboratory/timemachineimageanalysis) then manually reviewed these stitched images. This review yielded positions containing candidate barcode RNA FISH positive cells which we then re-imaged for verification at 60X magnification in multiple Z-planes. If we were uncertain about the fluorescence signal in a candidate cell (e.g., abnormal localization pattern, non-specific signal in multiple channels), we excluded the cell from imaging during subsequent rounds of RNA FISH or immunofluorescence. For quantification of RNA FISH images we used custom MATLAB software available at: https://github.com/arjunrajlaboratory/rajlabimagetools. Briefly, the image analysis pipeline includes manual segmentation of cell boundaries, thresholding of each fluorescence channel in each cell to identify individual RNA FISH spots, and then extraction of spot counts for all channels and cells. Notably, for some genes, we were not able to quantify expression in a few cells because of grossly abnormal or non-specific fluorescence signal (i.e. schmutz) or because we lost a cell during sequential hybridizations. We excluded data from these cells from analyses and as a result, some plots may contain slightly different numbers of points for different genes. For quantification of cell numbers for determining proliferation rate, we used custom MATLAB software available at https://github.com/arjunrajlaboratory/colonycounting_v2. Briefly, the image analysis pipeline involves stitching the tiled DAPI images, identifying individual cells based on DAPI signal, and then extracting cell counts from the entire well.

### Flow sorting of cells based on cycling speed

To sort hiF-T cells by cycling speed, we labeled hiF-Ts with CellTrace Yellow (Invitrogen #C34567) at a final concentration of 10uM. Briefly, we dissociated the cells, centrifuged and resuspended in 1XDPBS for two washes, added CellTrace Yellow to the cells in suspension, and incubated at 37°C for 30 minutes. After incubation, we resuspended in five volumes of media to remove free dye, centrifuged and resuspended in fresh media, and replated onto 10 cm plates. For experiments with barcoded hiF-Ts, we stained hiF-T cells with CellTrace Yellow immediately before performing lentiviral transduction of clone barcodes. Generally, cells were harvested for FACS sorting 3-4 days following labeling. Fast and slow cycling cells were determined by examining the distribution of CellTrace Yellow signal during FACS sorting, corresponding to the dimmest 15% of cells and brightest 15% of cells respectively.

### CRISPR knockdown construct lentivirus generation

To knockdown expression of several priming markers identified by Rewind, we generated lentiviral CRISPR constructs. The hiF-T cells are sensitive to seeding density and often do not survive at low densities, making clonal bottlenecking difficult. Hence, because we were unable to confirm sample-wide genetic indels via precise selection of knockout clones, we refer to the effect here as knockdown instead of knockout despite using CRISPR/Cas9. To design each CRISPR knockdown construct, we selected 2-4 guides per gene from a genome-wide database designed using optimized metrics in (Doench et al., 2016), generally prioritizing guides with a higher Rule Set 2 score when possible. We ordered pairs of complementary forward and reverse single stranded oligonucleotides (IDT) for each guide containing compatible overhang sites for insertion into a lentiCRISPRv2-blast (Addgene #83480) backbone, which simultaneously encodes Cas9 and an insertable target guide DNA (gDNA). We resuspended each oligo to 25 uM in NF-H2O and performed phosphorylation and annealing of each oligo pair by combining the oligos with 1X T4 ligase buffer (NEB #B0202S) and polynucleotide kinase (NEB #M0201S) and incubating in a thermocycler (Veriti #4375305) with the following settings: 37°C for 30 min, 95°C for 5 min, and ramping down to 25°C at 5°C/min. Then, we inserted our annealed oligos into the lentiCRISPRv2-blast backbone via Golden Gate assembly; we combined 50 ng of backbone with 25 ng of annealed oligo along with T4 ligase buffer (NEB #B0202S), T4 ligase (#M0202S), and BsmBI restriction enzyme (NEB #R0739S) and incubated in a thermocycler (Veriti #4375305) with the following settings: 50 cycles of 42°C for 5 min and 16°C for 5 min followed by 65°C for 10 min. To grow out plasmid before packing, we performed heat-shock transformation at 42°C in Stbl3 *E. coli* cells for each guide before plating on LB plates with ampicillin and incubating at 225 rpm and 37°C for 8-12 hours. Individual colonies were picked for each guide and grown out in 5 mL liquid cultures of LB with ampicillin. Then, we pelleted each liquid culture by centrifugation and used the Monarch Plasmid Miniprep Kit (NEB #T1010L) to isolate plasmid according to the manufacturer’s protocol. Guides were verified by Sanger sequencing of the isolated plasmid.

### Lentivirus packaging of CRISPR knockdown constructs

To package sequence-verified plasmids for each guide into lentivirus, we first grew HEK293T to 65%-80% confluency in 6-well plates in DMEM + 10% FBS without antibiotics and on 0.1% gelatin (#ES-006-B). For each individual plasmid, we combined 0.5 ug of pMD2.G, 0.883 ug of pMDLg, 0.333 ug of pRSV-Rev, and 1.333 ug of the plasmid in 200 uL of Opti-MEM. After vortexing, we added 9.09 uL of polyethylenimine (Polysciences #23966) and incubated for 15 min at room temperature before adding the final mixture for each plasmid to a HEK293-containing well of the 6-well plates. After 4-6 hours, we aspirated the media and added fresh hiF-T GM. After 48 hours, we filtered the virus-laden media through a 0.45µm PES filter (Millipore-Sigma #SE1M003M00) and stored 1.5 mL aliquots in cryovials at -80ºC.

### Transduction with lentiviral CRISPR knockdown constructs

To transduce hiF-T cells, we freshly thawed virus-laden media on ice, added it to dissociated cells with 4 ug/mL of polybrene (Millipore-Sigma #TR-1003-G), and plated 50,000 cells/well in a 6-well plate coated with Attachment Factor (Fisher #S006100). We used 400 uL/well of virus-laden media, aiming for a high MOI. After plating hiF-T cells with virus, we performed a 30 min incubation at room temperature before centrifuging the 6-well plate at 930g for 30 min at room temperature. After 24 hours, we passaged the cells from 2 wells onto 10 cm plates and began selection in 2.5 ug/mL of blasticidin. This selection was removed after 7 days, which corresponded with when cell death was near complete in control hiF-Ts we cultured in parallel containing no lentiviral constructs. We wanted to minimize the effect of blasticidin on reprogramming, so we cultured the cells for 7 days without blasticidin and without puromycin before reprogramming via OKSM induction.

### Immunoblotting analyses of whole-cell lysates

For immunoblotting analysis, we prepared whole-cell lysates without sonication as described in (Dou et al., 2017). Briefly, cells were lysed in buffer containing 20mM Tris, pH 7.5, 137 mM NaCl, 1 mM MgCl2, 1mM CaCl2, and 1% NP-40 supplemented with 1:100 Halt protease and phosphatase inhibitor cocktail (Thermo Fisher #78430) and benzonase (Novagen #707463) at 12.5 U/mL. The lysates were rotated at 4°C for 30min and boiled at 95°C in the presence of 1% SDS. The resulting supernatants after centrifugation were quantified by using the Pierce Rapid Gold PCA Protein Assay Kit (Thermo #A53225) and equal amounts were subjected to electrophoresis using NuPAGE 4%-12% Bis-Tris precast gels (Thermo #NP0335BOX). Afterwards, we cut the nitrocellulose membrane into separate strips for each protein being probed, and we used 5% milk in TBS supplemented with 0.1% Tween20 (TBST) to block the membrane at room temperature for 30 min. Primary antibodies were diluted in 5% milk in TBST and incubated at 4°C overnight. The membrane was washed 3 times with TBST, each for 10 min, followed by incubation of secondary antibodies at room temperature for 1 hour in 5% milk in TBST. The membrane was washed again 3 times and imaged on an Amersham Imager 600 (GE Healthcare). The primary antibodies used were 1:1000 rabbit anti-SPP1 (Proteintech #22952-1-AP), 1:1000 rabbit anti-LSD1 (Cell Signaling Technology #2184S), 1:1000 rabbit anti-beta actin (Cell Signaling Technology #4970S), and 1:2000 rabbit anti-histone H3 (Abcam #AB1791). The secondary antibody used was 1:5000 HRP goat anti-rabbit (Bio-Rad #1706515).

### Bulk RNA-sequencing and analyses

We conducted standard bulk, paired-end (37:8:8:38) RNA-sequencing using a RNeasy Micro kit (Qiagen #74004) for RNA extraction, NEBNext Poly(A) mRNA Magnetic Isolation Module (NEB #E7490L), NEBNext Ultra II RNA Library Prep Kit for Illumina (NEB #E7770L), NEBNext Multiplex Oligos for Illumina (Dual Index Primers Set 1) oligos (NEB EE7600S), and an Illumina NextSeq 550 75-cycle high-output kit (Illumina #20024906), as previously described (Mellis et al., 2021). As previously described, we aligned RNA-seq reads to the human genome (hg38) with STAR and counted uniquely mapping reads with HTSeq (Dobin et al., 2013) and outputs count matrix. The counts matrix was used to obtain transcripts per million (TPM) and other normalized values for each gene using scripts provided at: https://github.com/arjunrajlaboratory/RajLabSeqTools/tree/master/LocalComputerScripts. We performed differential expression analysis in R using DESeq2and with data from at least 2 biological replicates for each sample and condition.

### Transcription factor overrepresentation and binding motif analysis

We used ChEA3 (Keenan et al., 2019) to identify possible upstream regulators of a subset of the positive and negative priming markers. ChEA3 integrates data about associations between transcription factors and target genes from multiple assays and genomic analyses. We used the jsonlite package in R to access the ChEA3 application programming interface (API) to submit queries to perform transcription factor target overrepresentation analysis. We submitted separate queries for the best 50 negative priming markers (i.e., lowest log2(fold change) for primed cells / nonprimed cells) and for the best 50 positive priming markers (i.e., highest log2 (fold change) for primed cells / nonprimed cells). We sorted the output from each query by integrated mean rank across each of ChEA3’s databases. For several putative upstream regulators for the negative priming markers (*TWIST2, PRRX2, OSR1*, and *SNAI2*), we extracted binding motifs from the JASPAR database (Castro-Mondragon et al., 2022) using the monaLisa package in R. Then, we used the scanMotifGenomeWide.pl command within HOMER (Heinz et al., 2010) to look for instances of each motif across the genome and output a text file with the results. To simultaneously visualize binding motifs for each of these transcription factors upstream of SPP1 and FTH1, we combined the text files for each motif into a single BED file in R and uploaded it as a custom track in the Integrated Genome Viewer (Robinson et al., 2011).

### Calculating odds ratios for priming given expression of priming markers

To measure the relative explanatory power of cycling speed and a subset of our identified priming markers, we calculated log(odds ratios). For cycling speed (dichotomous variable), we asked if a cell is in a given cycling speed group, what are its odds of being primed. The odds ratio for each cycling speed was calculated with the corresponding standard error using the single-cell RNA-sequencing dataset generated in Figure 3A. For each primed marker (continuous variable), we asked if a cell has high expression (in the top 10% of expression values per cell) of a given marker, what are its odds of being primed. Briefly, we built 2 × 2 contingency tables in each case and calculated the log(odds ratio) as (odds of being primed with high expression of gene x) / (odds of being primed without high expression of gene x). The standard error of the log(odds ratio) was approximated by the square root of the sum of the reciprocals of the four frequencies (Bland & Altman, 2000). The odds ratios for each marker were calculated with corresponding standard error separately in n = 3 biologically independent single-cell RNA-sequencing datasets identifying primed and nonprimed cells in the initial population and aggregated using a random-effects model using the metafor package in R. Additionally, we generated logistic regression models to evaluate the contributions of cycling speed, expression of our identified priming markers, and the interactions between these terms in predicting priming. For each primed marker, we generated a separate logistic regression model using the glm command in R and generated a p-value for each term coefficient determined by the likelihood ratio test. For cycling speed as well as each primed marker individually, we generated single-term logistic regression models (i.e., priming = B0 + B1*cycling speed or priming = B0 + B1*gene expression) and calculated R^2^ values to calculate the percent variation in priming explained by each priming marker or by cycling speed alone.

### Mixing coefficient calculation and nearest neighbor analyses

We used a quantifiable approach to measure the gene expression relatedness of different barcoded clones as described in (Goyal et al., 2021). For each pair of barcoded clones, we calculated the nearest neighbors for each cell in the 50-dimensional principal component space. We then classified the neighbors as “self” if the neighbors were from the same barcode clone or “non-self” if they belonged to the other barcode clone. We calculated the mixing coefficient as follows: mixing coefficient = (number of non-self neighbors) / (number of self-neighbors). A mixing coefficient of 1 would indicate perfect mixing such that each cell has the same number of self and non-self neighbors. A mixing coefficient of 0 would indicate that there is no mixing and that each cell within a barcoded clone lies far away from the other barcoded clone in the principal component space. All cases in which the calculated mixing coefficient was ≥1 were considered perfect mixing. The higher the mixing coefficient, the higher the transcriptional relatedness of the barcoded clones analyzed. As the number of nearest neighbors depends on the size (number of cells) of a clone, we performed this analysis between cells of similar clone size. We performed this analysis only on clones with at least 4 cells in each split.

### Cluster probability distribution and Jensen-Shannon distance analyses

To more sensitively measure the gene expression relatedness of different barcoded clones, we calculated Seurat cluster probability distributions and Jensen-Shannon distance as in (Jiang et al., 2022). For each clone barcode within a dataset, we found how its associated cells partitioned across Seurat clusters. We then divided the raw number of cells per cluster by the total number of cells found in that cluster within that dataset, then normalized all cluster proportions to sum to 1 to get “probability distributions” for each barcoded clone. To generate the probability distributions we might expect from random chance, we averaged the results from the normalization as above for 1000 random samples of a matched number of barcoded cells, noting that for all barcodes this distribution was approximately uniform. We calculated the Jensen-Shannon distance between our observed barcode probability distribution and the averaged random probability distribution largely as previously described in (Arumugam et al., 2011), one exception being that we chose to use log base 2 to calculate the Kullback-Leibler divergence so that maximally different samples would have a Jensen-Shannon distance of 1. To determine the significance of our calculated Jensen-Shannon distance, we took another 1000 random samples of a matched number of cells, and for the probability distribution associated with each random sample we calculated the Jensen-Shannon distance from the averaged random probability distribution. We performed this analysis only on clones with at least 4 cells in each split.

### Data and Code Availability

All raw and processed data as well as code for analyses in this manuscript can be found at https://www.dropbox.com/sh/ulu6728tcp49dv2/AAAPwLYQiVLloH_JL38lvTj6a?dl=0.

## Author Contributions

NJ and AR conceived and designed the project with valuable input from YG, IAM, and BE. NJ designed, performed, and analyzed all experiments, supervised by AR. IAM assisted NJ with optimizing reprogramming conditions in hiF-T cells. YG, IPD, SR, CLJ, and BE helped with some hiF-T cell maintenance and reprogramming experiments. IPD assisted NJ with automated RNA FISH and DAPI scans. MD and CC assisted NJ with generating RNA FISH probes and single-blind, manual quantification of RNA FISH counts and iPSC colony counts. SR assisted NJ with generating CRISPR knockdown constructs. The protocol for barcode library generation and subsequent initial pipeline for retrieval of barcodes from genomic DNA-sequencing were developed by BE. The initial pipeline for retrieval of barcodes from copy DNA and matching single-cell sequencing data was developed by YG. The analyses and scripts for measuring similaring in single-cell RNA gene expression profiles were conceived and developed by YG and CLJ. NJ and AR wrote the manuscript.

The authors read and approved the final manuscript.

## Acknowledgements

We thank members of the Arjun Raj Lab, particularly Lauren Beck, Phil Burnham, Lee Richman, Yael Heyman, Karun Kiani, Eric Sanford, Allison Cote, and Cat Triandafillou for insightful discussions related to this work. We thank Blake Caldwell of the Marisa Bartolomei lab, Jingchao Zhang of the Ken Zaret lab, Chris Lengner, as well as Wenli Yang and Rachel Truitt of the Penn iPSC core for advice on iPSC handling and reprogramming. We thank Daniel Xu of the Shelley Berger lab for assistance with immunoblotting protocols. We thank the Genomics Facility at the Wistar Institute, especially Sonali Majumdar and Sandy Widura for assistance with single-cell partitioning and addition of 10x cell identifiers. We thank the Flow Cytometry Core Laboratory at the Children’s Hospital of Philadelphia Research Institute for assistance with flow cytometry and fluorescence-activated sorting. Finally, NJ thanks his beloved cat Brioche for keeping him company on her favorite chair or in her favorite box during hours of writing and proofreading (and for some unanticipated edits).

NJ acknowledges support from the Michael Brown Fellowship, NIH T32 GM717044, and NIH F30 HD103378. YG acknowledges support from the Burroughs Welcome Fund Career Awards at the Scientific Interface, a grant from Research Catalyst Program from the McCormick School of Engineering at Northwestern University, and Northwestern University’s startup funds. IAM acknowledges support from NIH F30 NS100595. BE acknowledges support from NIH F30 CA236129, NIH T32 GM007170, NIH T32 HG000046. CLJ acknowledges support from NIH F30 HG010822, NIH T32 DK007780, and NIH T32 GM007170. AR acknowledge support from TR01 GM137425, NIH R01 CA238237, NIH R01 CA232256, NIH 4DN U01 DK127405, and NSF EFRI EFMA19-33400.

