## Supplementary Figures for "Retrospective identification of intrinsic factors that mark pluripotency potential in rare somatic cells"

### Supplementary Figure 1

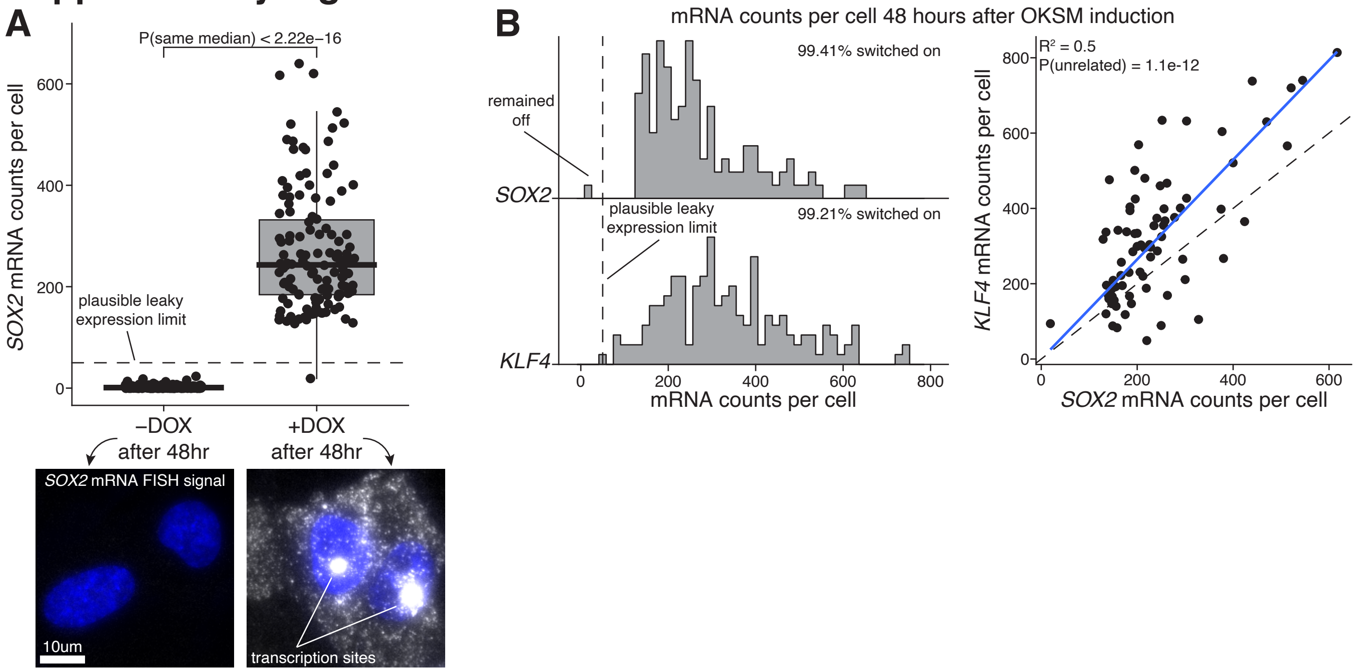

### Supplementary Figure 2

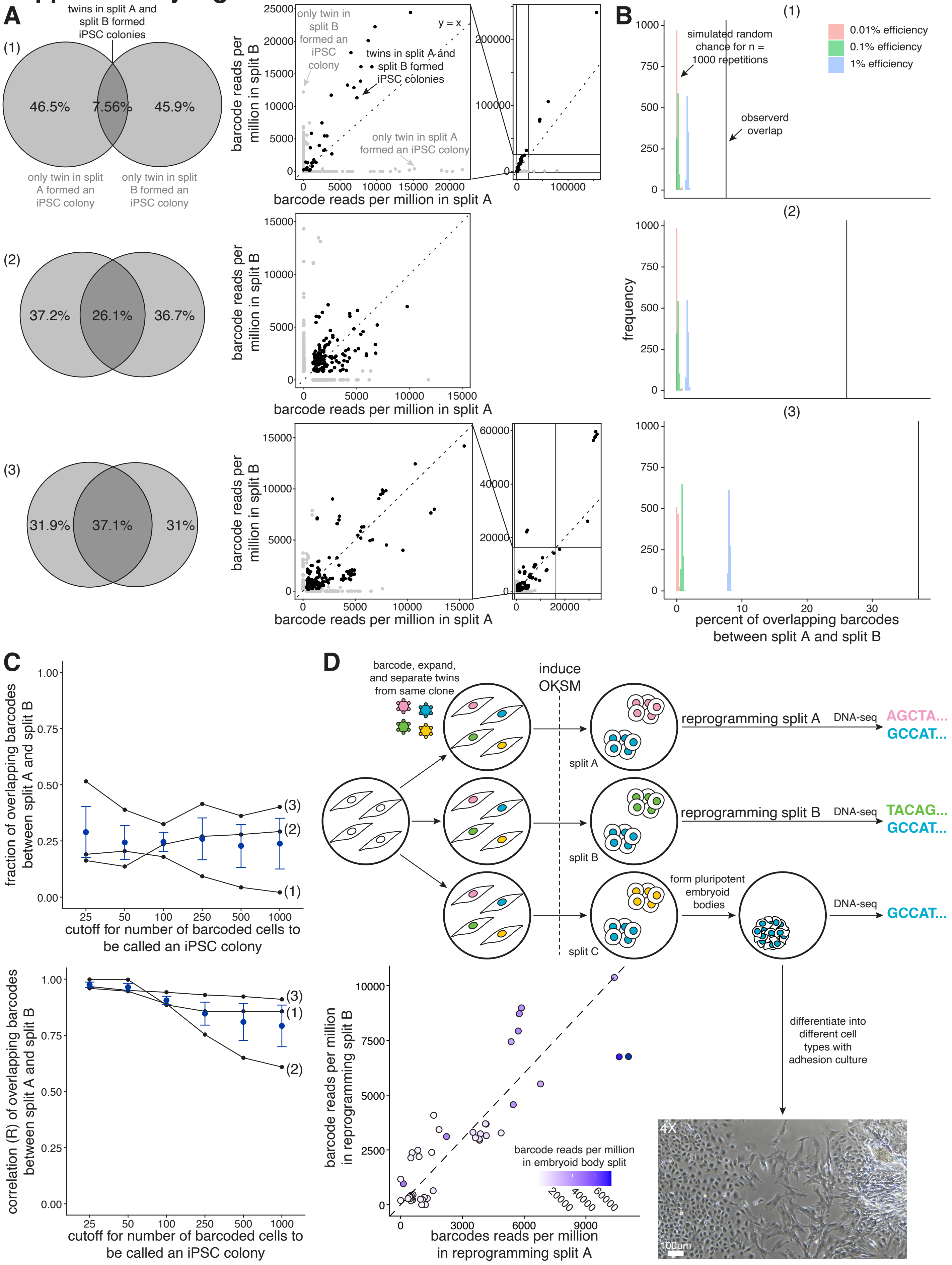

### Supplementary Figure 3

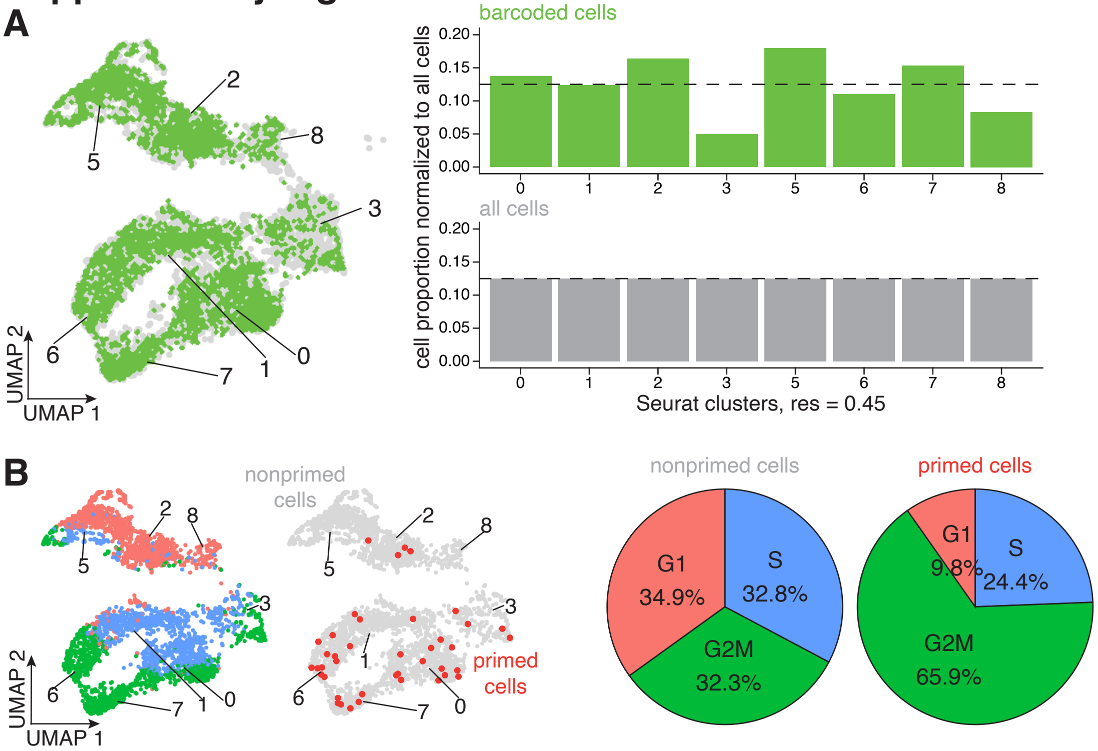

### Supplementary Figure 4

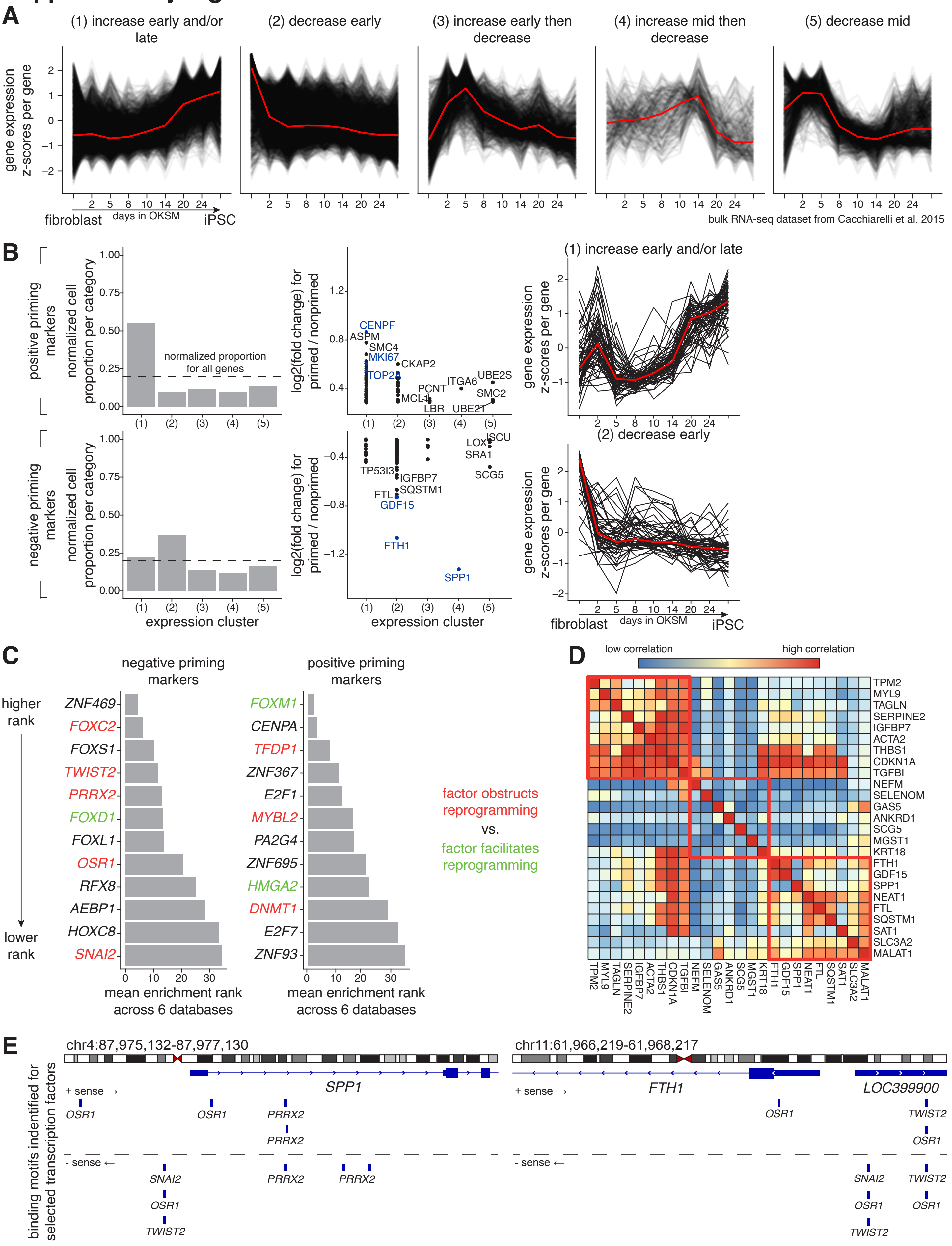

### Supplementary Figure 5

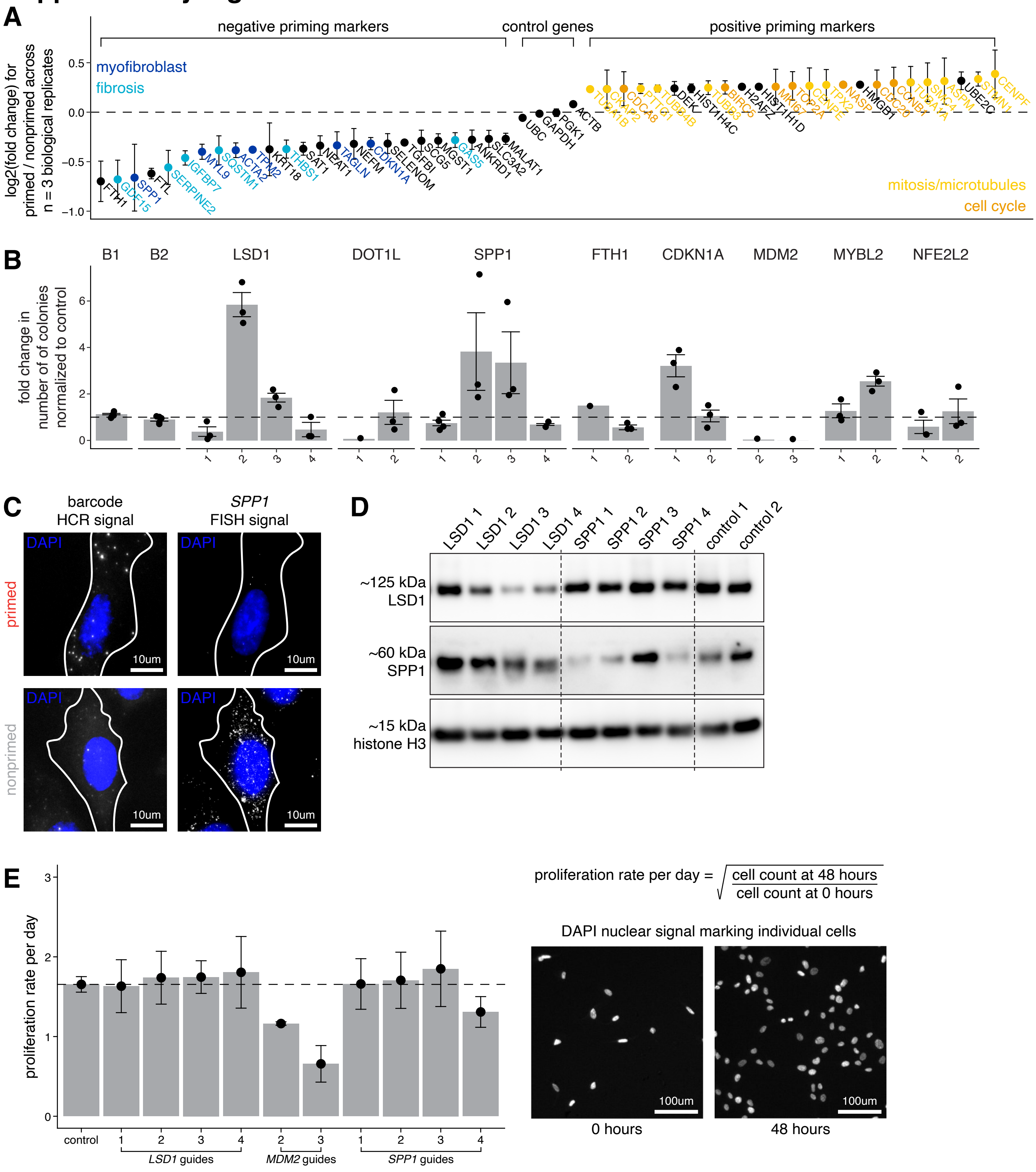

### Supplementary Figure 7

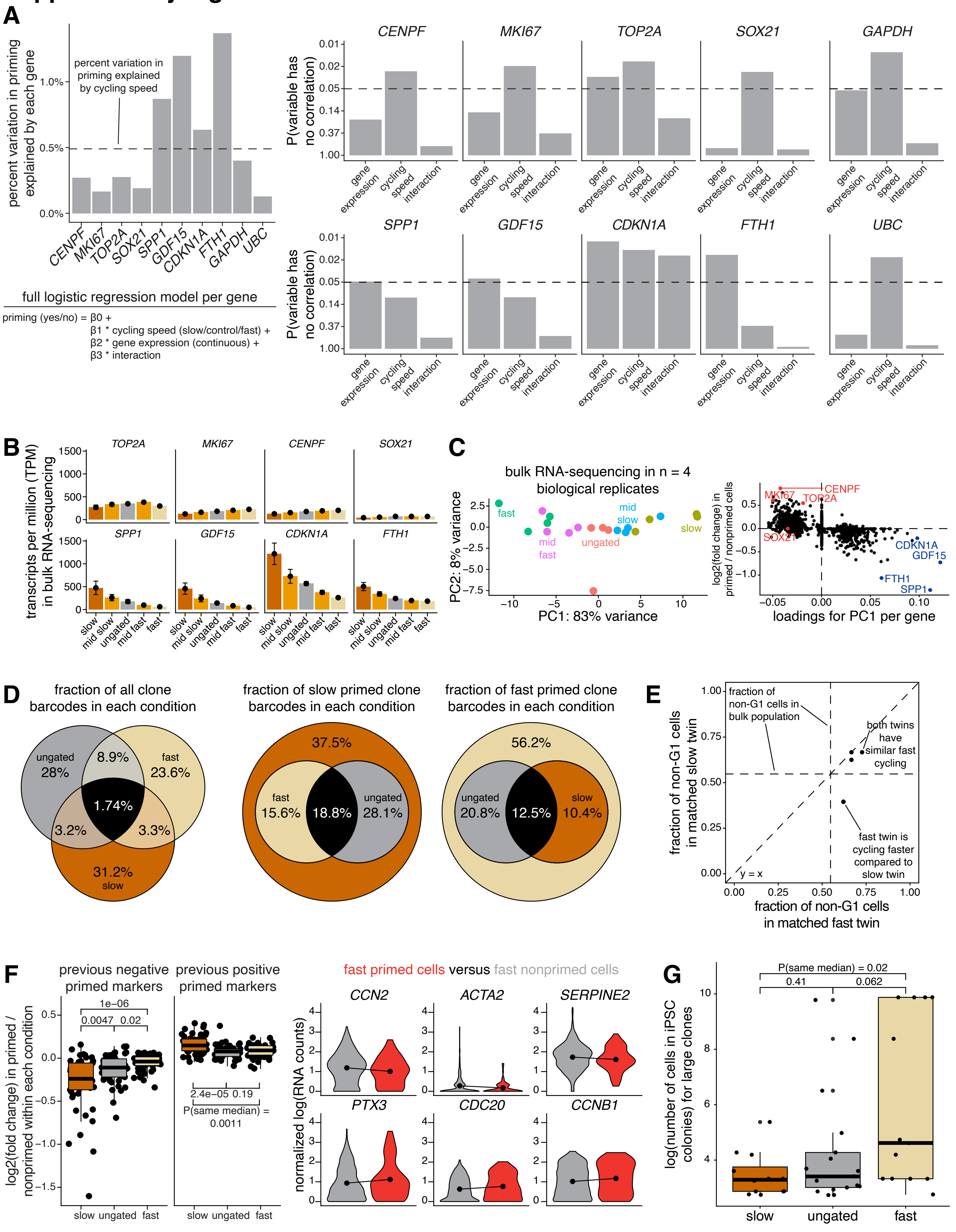

### Supplementary Figure 6

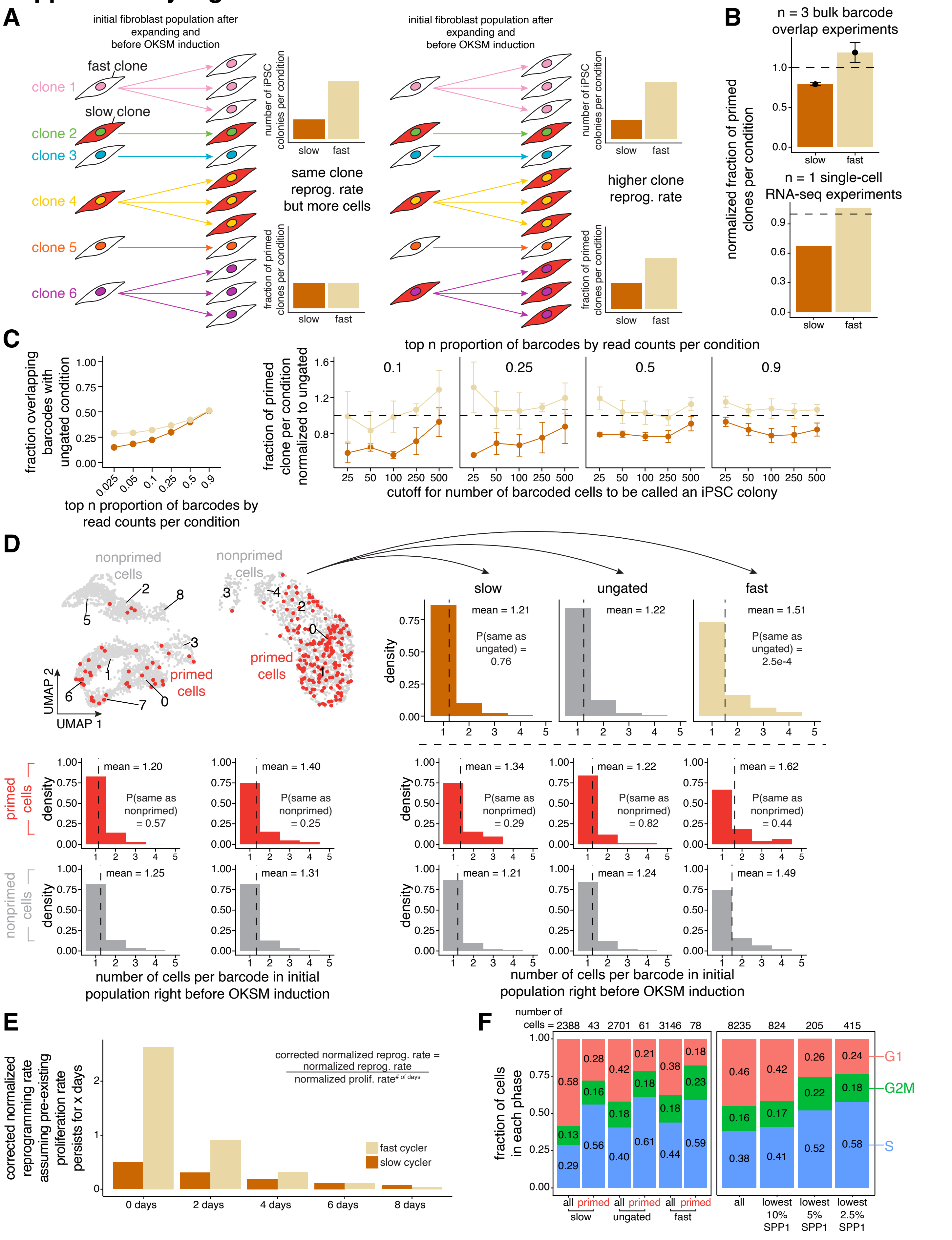

### Supplementary Figure 8

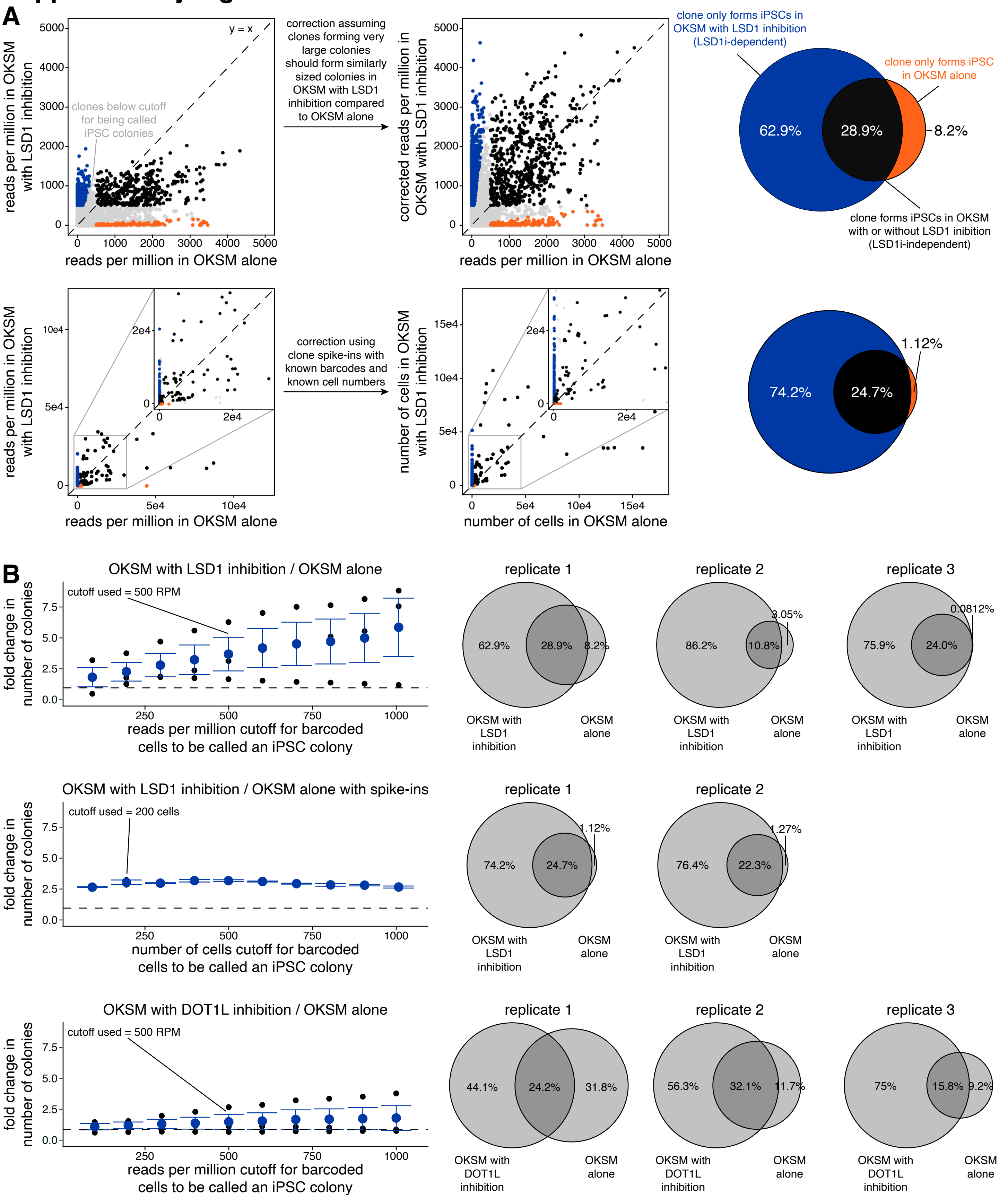

### Supplementary Figure 9

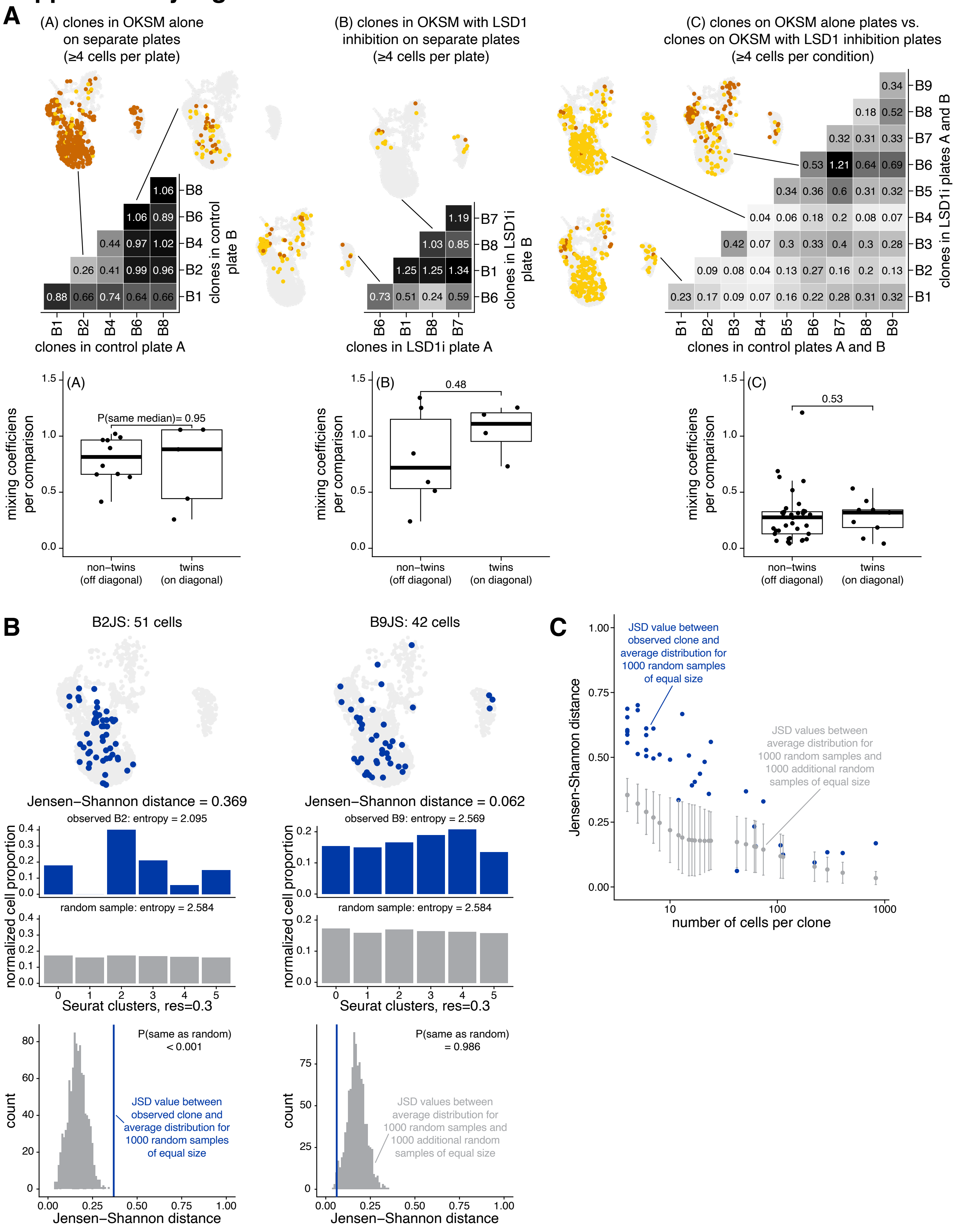

### Supplementary Figure 10

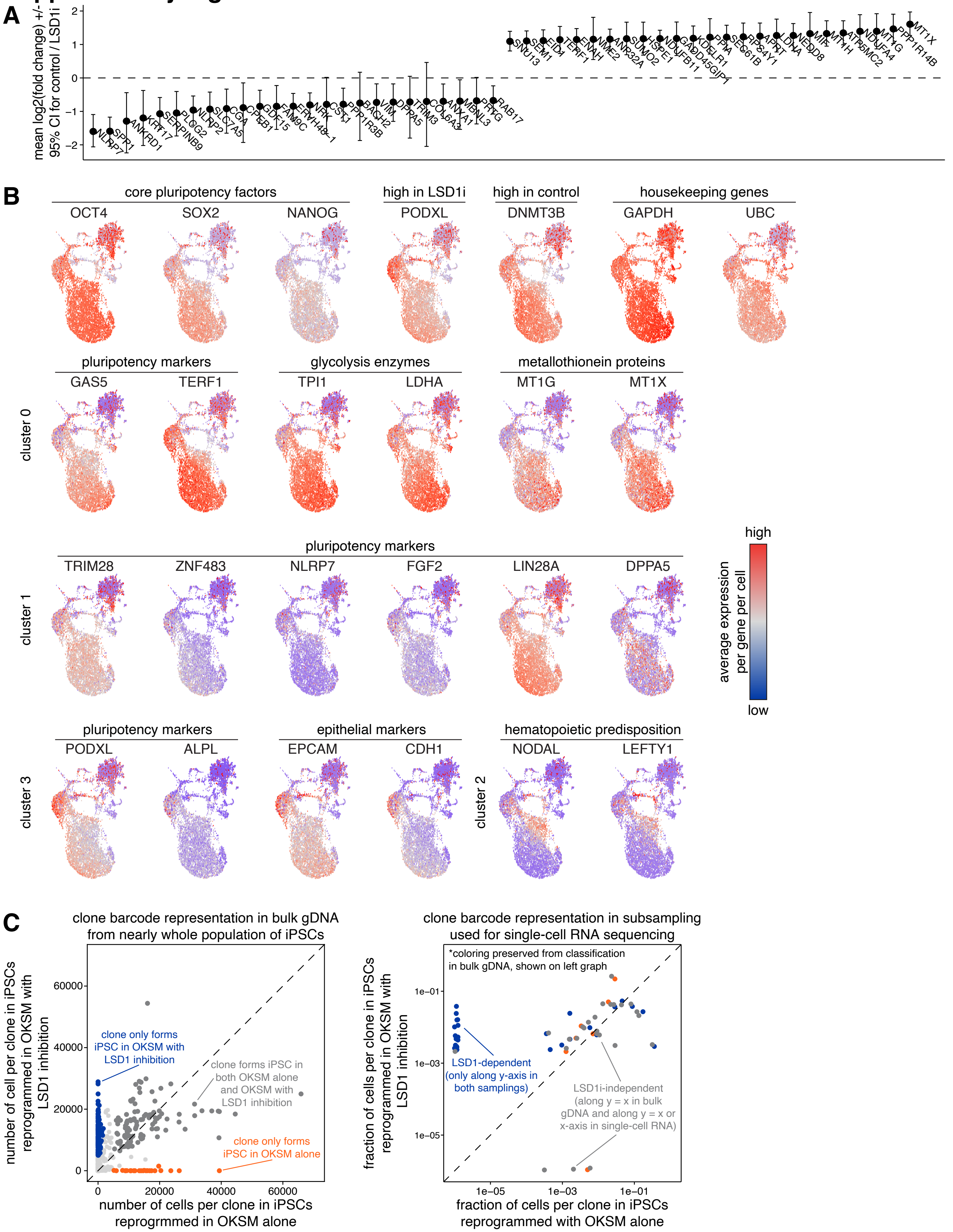

#### Supplementary Figure Captions

**Figure S1: Measurement of OKSM induction with single-cell resolution in hiF-T cells**

A. We measured *SOX2* expression levels by FISH in individual hiF-T cells without and with OKSM induction after 48 hours in culture. Each dot represents an individual cell. Shown are representative single-molecule FISH images contrasted equally with DAPI in blue and *SOX2* mRNA FISH signal in white. Large, high intensity nuclear spots likely represent sites of active transcription. In uninduced hiF-Ts we observed a maximum SOX2 expression level of 31 counts, so we set a plausible limit for leaky OKSM expression of 50 counts (i.e., high expression of OKSM without induction by adding doxycycline).

B. We wanted to verify that most hiF-Ts switched on expression of OKSM following induction. We measured *SOX2* and *KLF4* expression levels by FISH in individual hiF-Ts 48 hours after OKSM induction. We classified cells with expression levels above a maximum level seen in uninduced cells (i.e., "plausible leaky expression limit") as being "on" and cells with expression levels below this limit as being "off". To verify if individual OKSM factor induction levels were relatively stoichiometric as would be expected with the polycistronic cassette, we plotted *SOX2* mRNA counts versus *KLF4* mRNA counts per individual cells and fit a linear regression model in R.

**Figure S2: Quantification of clone barcode overlap in Rewind splits demonstrates priming has robust memory across cell division**

A. For each clone barcode identified by sequencing genomic DNA in reprogrammed iPCS, we plotted its abundance in the corresponding split A (x-axis) and split B (y-axis). (1), (2), and (3) represent n = 3 independent biological replicates. Clone barcodes present in both split A and split B are colored in black as points and in gray on the corresponding Venn diagram, while barcodes present only in split A or split B are colored light gray as both points and on the corresponding Venn diagram. For (1) and (3), we plotted pullout plots because of the presence of clones with extremely high numbers of reads.

B. We compared the observed barcode overlap in each experiment to the distribution of 1000 simulations assuming a "no memory" model wherein cells becoming iPSCs were chosen at random in each split (see Methods). We generated null distributions for 1000 simulations using one of several reprogramming frequencies (0.01% in light red, 0.1% in light green, and 1% in light blue).

C. For each experiment in Figure 1D, we used a cutoff of 250 reads per million to classify cells as iPSC colonies. We measured barcode overlap across different cutoff values. We also measured the correlation for each clone barcode between its abundance in split A and its abundance in split B by fitting a linear regression model in R including the origin. Again, we measured the correlation across different cutoff values.

D. We transduced hiF-T cells at an MOI of ~1 with our barcode library. After 4 cell divisions, we divided the culture into splits (A, B, and C). In all splits, we reprogrammed cells into iPSCs via induction of OKSM. In splits A and B we performed barcode DNA-sequencing on the resulting iPSCs to quantify clone abundance in each split. In split C, we generated embryoid bodies from the resulting iPSCs in suspension culture and performed barcode DNA-sequencing on the resulting embryoid bodies to quantify clone abundance. We plotted clone abundance in split A (x-axis), split B (y-axis), and split C (z-axis, represented by color). We plated a small amount of embryoid bodies from split C in adhesion culture. The embryoid bodies differentiated and gave rise to cell types of varying morphology, as shown in this representative light microscopy image.

**Figure S3: Distribution of barcoded cells and cell cycle phases in UMAP clusters**

A. We measured the distribution of barcoded cells across clusters while maintaining the organization provided by UMAP in our Rewind experiment in Figure 1F. Barcoded cells here are all cells in which we can conclusively associate a single clone barcode, and not GFP positive cells.

B. We classified cells in our single-cell RNA-sequencing dataset by cell cycle phase based on Seurat's CellCycleScoring command, which makes use of phase-specific markers as described in (Tirosh et al., 2016). While maintaining the organization provided by UMAP in our Rewind experiment in Figure 1F, we recolored each cell by its assigned cell cycle phase in the left UMAP and by its priming status in the right UMAP. Cells in G1 are light red, in S are blue, and in G2M are green. For primed and nonprimed cells, we measured the relative proportion of cells in each cell cycle phase and plotted the corresponding pie charts.

**Figure S4: Positive and negative priming markers are associated with distinct gene expression patterns during reprogramming and genes in each category may be coordinated by common upstream regulators**

A. We performed hierarchical clustering using the hclust package in R to categorize genes based on broad expression patterns when reprogramming from hiF-T cells into iPSCs using bulk RNA-sequencing data from (Cacchiarelli et al., 2015). To normalize for genes with different expression level ranges, we calculated a z-score (z-score = (observed expression value - average expression value) / (standard deviation)) of gene expression for each timepoint per gene. Each line in black is the expression pattern for an individual gene, while each line in red is the average expression pattern for all genes in that category.

B. We used the same hierarchical clustering method to categorize the best 100 positive priming markers (i.e., highest log2(fold change) between primed and nonprimed cells) and the best 100 negative priming markers (i.e., lowest log2(fold change) between primed and nonprimed cells). For each set of priming markers, we plotted the proportion of genes in each category normalized to the number of genes in each category. To better visualize how the best markers in each set were categorized, we also plotted the distribution of log2(fold change) values per gene in each category.

C. We used ChEA3 (Keenan et al., 2019) to identify possible upstream regulators of the positive and negative priming markers. Here, a higher rank corresponds to a stronger putative regulator. Identified regulators are colored in red if previous studies demonstrated a role in obstructing reprogramming and are colored in green if previous studies demonstrated a role in facilitating reprogramming.

D. We asked whether sets of negative priming markers might have coordinated expression. For the best 25 negative priming markers (i.e., lowest log2(fold change)), we calculated pairwise values for how often each gene is co-regulated with every other gene for each of the best 200 upstream regulators (i.e., lowest mean rank) and normalized those values by number of occurrences per gene to get a correlation matrix. We identified 3 broad sets of genes that could have coordinated regulation, outlined here by red squares.

E. We identified binding motifs for *TWIST2*, *PRRX2*, *OSR1*, and *SNAI2* across the genome (see Methods). We visualized these binding motifs in the Integrated Genome Viewer (Robinson et al., 2011). Here, we show the presence of binding motifs for multiple queried transcription factors upstream of *SPP1* and *FTH1*, two prominent negative priming markers.

**Figure S5: Validation of identified priming markers in additional Rewind experiments and by CRISPR knockdown**

A. We performed n = 3 independent biological Rewind experiments and calculated average log2(fold change) values for each gene in primed cells versus nonprimed cells. Factors associated with myofibroblasts are in dark blue, with activated fibroblasts are in light blue, with cell cycle are in dark yellow, and with mitosis are in light yellow. Metric shown is mean +/- standard error.

B. We designed CRISPR guides to knockdown expression of several priming markers. We quantified the number of iPSC colonies formed for each guide (labeled numerically) for each gene and reported each measurement as a normalized value relative to a backbone control lacking a target guide RNA (i.e, B1 or B2). Metric shown is mean +/- standard error for 2-3 technical replicates per guide.

C. We performed single-molecule RNA FISH to measure gene expression of *SPP1* in primed and nonprimed cells in Figure 2B. Shown are representative images. The first column has clone barcode RNA FISH signal in white and DAPI in blue. The second column has *SPP1* RNA FISH signal in white and DAPI in blue.

D. We performed immunoblotting to see if mRNA knockdown correlated with protein knockdown with our CRISPR guides for *LSD1* and *SPP1*. This blot represents lysates run on the same gel and membrane, but the

membrane was cut into 3 sections before staining with primary and secondary antibody. We stained for LSD1, SPP1, and histone H3 as a loading control.
E. To measure how knockdown for each guide affected proliferation rate, we measured cell number for each guide at 0 hours and 48 hours after fixing and staining for DAPI. We quantified the proliferation rate per day per guide as proliferation rate per day = (cell count at 48 hours / cell count at 0 hours)^1/2. The dashed line is the proliferation rate per day for backbone control lacking a target guide RNA (i.e., “control’). Metric shown is mean +/- standard error for n = 2 biological replicates.

**Figure S6: Fast cycling cells have an intrinsically higher rate of reprogramming beyond simply having an increased number of cells entering the reprogramming process**
A. The increased number of colonies formed from fast cycling cells observed could be simply because each fast primed clone results in more iPSC colonies via an increased number of primed progeny entering reprogramming (i.e., “same clone reprog. rate but more cells”), or it could be because a higher fraction of fast cycling clones are primed (i.e., “higher clone reprog. rate”).
B. To distinguish the possibilities in Figure S6A, we performed a similar barcode overlap experiment as in Figure 1B but with split B we sorted cells on cycling speed and performed barcode DNA-sequencing before inducing OKSM. After reprogramming split A into iPSCs via OKSM induction, we measured the fraction of clones in each cycling speed group from split B forming iPSCs in split A. Metric shown is mean +/- standard error for n = 3 independent biological replicates. We also performed a similar analysis using our single-cell RNA-sequencing dataset from Figure 3.
C. For our analysis in Figure S6B, we measured the amount of barcode overlap between slow, ungated, and fast fibroblasts from our cycling speed sort populations. We measured this barcode overlap with different cutoffs for the proportion of barcodes to include in each condition to filter out poorly represented barcodes that could represent PCR amplification or sequencing artifacts. Metric shown is mean for n = 3 independent biological replicates. Additionally, we measured the normalized fraction of clones per condition across different barcode proportion cutoffs as well as different cutoffs for the minimum number of cells (determined by spike-in controls of known clone barcodes) to be called an iPSC colony. Metric shown is mean +/- standard error for n = 3 independent biological replicates. Fast cyclers are in khaki, slow cyclers are in light brown, and ungated cells were used for normalization and are indicated with the dotted line. For Figure S6B, we are showing results using a colony cutoff of 25 and a barcode proportion cutoff of 0.5.
D. To determine if primed clones reprogram at higher efficiency by virtue of having more cells at the start of OKSM induction, we measured the number of starting cells per clone in our single-cell RNA-sequencing datasets from Figure 1 and from Figure 3. The left set of graphs show the distribution of number of starting cells per clone for primed (in red) and nonprimed (in gray) clones in bulk for n = 2 independent biological replicates. For the second replicate we could further classify cells by cycling speed, which is shown on the right set of graphs. We compared distributions for bulk cycling speed categories and between primed and nonprimed cells within each cycling speed category. The dotted vertical line is the mean for each distribution. P-values comparing means were calculated using the Student’s t-test.
E. To measure if differences in growth rate alone could explain the observed differences in iPSC generation between fast cyclers and slow cyclers, we calculated corrected reprogramming efficiencies assuming pre-existing differences in proliferation rate persisted at increasingly later time points after OKSM induction as an extreme model. To do so, we calculated corrected reprogramming efficiency = (normalized reprogramming rate in Figure 2E) / (normalized proliferation rate in Figure 2D)^(number of days).
F. We classified cells in our single-cell RNA-sequencing data set by cell cycle phase based on Seurat’s CellCycleScoring command, which makes use of phase-specific markers as described in (Tirosh et al., 2016). Here, G1 is in light red, G2M is in green, and S is in blue. We plotted the fraction of cells in each phase for all cells versus primed cells within each cycling speed. Additionally, we plotted the fraction of cells in each phase for all cells versus different subsets of cells with increasingly lower SPP1 expression (i.e., cells with the lowest 10%, 5%, and 2.5% of SPP1 expression values). The number of cells in each category are indicated.

**Figure S7: Explanatory power and expression level analysis of select priming markers across cycling speed**

A. We generated logistic regression models to evaluate the contributions of cycling speed, expression of our identified priming markers, and the interactions between these terms in predicting priming using our single-cell RNA-sequencing dataset from Figure 3. For each primed marker, we generated a separate logistic regression model using the glm command in R and plotted the p-value (on an inverse y-axis scale) for each term coefficient determined by the likelihood ratio test. Additionally, for cycling speed as well as each primed marker we generated single term logistic regression models (i.e., priming = B0 + B1*cycling speed or priming = B0 + B1*gene expression) and calculated $R^2$ values to calculate the percent variation in priming explained by each individual term.

B. We performed bulk RNA-sequencing on populations of hiF-T cells sorted by cycling speed using the accumulation dye approach described in Figure 2C. We normalized read counts for each gene by calculating transcripts per million (TPM), and plotted the average TPM in each sorted population for a subset of our identified priming markers. Metric shown is mean +/- standard error for n = 4 independent biological replicates.

C. To visualize different axes of biological variability in our bulk RNA-sequencing dataset, we plotted expression profiles for each sample in principal component space. We extracted loadings for each principal component and plotted them for principal component 1 against the log2(fold change) values for each gene in primed versus nonprimed cells in our single-cell RNA-sequencing dataset from Figure 1. We marked a subset of positive priming markers in red and a subset of negative priming markers in blue.

D. In our single-cell RNA-sequencing dataset from Figure 3, we identified clone barcodes showing up in one or more accumulation dye sort populations and plotted the overlap as a Venn diagram. The percentages shown are for all barcodes in the dataset. We performed a similar analysis looking specifically at primed cells, asking what fraction of the slow primed barcodes show up in the ungated primed and fast primed populations and vice versa for fast primed cells.

E. We wondered whether clones showing up in both the fast primed and slow primed populations had similar cycling speeds. For 4 clones with ≥4 twins classified as fast primed and ≥4 twins classified as slow primed, we measured the fraction of cells not in G1 as an indicator of cycling speed. Each dot represents an individual barcode clone and dotted lines on each axis show the population fraction of cells not in G1 for all cells in the single-cell RNA-sequencing dataset for comparison. Clones near the y = x line indicate that the fast primed twins and slow primed twins have similar cycling speeds while clones in the bottom right quadrant indicate that the fast primed twins compared to the slow primed twins are cycling relatively faster.

F. Within each cycling speed population, we calculated log2(fold change) values for primed versus nonprimed cells using Seurat's FindMarkers command. We subsetted the list of log2(fold change) values for all genes to the top 25 negative priming markers on the left and the top 25 positive priming markers on the right and plotted the distributions for each cycling speed. P-values comparing sample means are calculated using the Wilcoxon rank sum test. To highlight what is different about fast primed (in red) versus fast nonprimed (in gray) cells, we chose a few of the most differentially expressed genes and plotted the distribution of log normalized RNA accounts using Seurat's VlnPlot command. Metric shown is mean with a connecting line segment for comparison.

G. We wondered if iPSC colonies arising from fast versus slow cycling primed cells had any phenotypic differences. In Figure 3H, we compared the distribution of iPSC colony size for each cycling speed across the whole range of size values. We plotted the same distributions for each cycling speed after subsetting for large iPSC colonies. We used a cutoff of 15 cells after normalizing read counts using our spike-in controls (see Methods). This cutoff, notably, is only a relative cell count because when harvesting iPSCs and performing library preparation we are subsampling the whole population of reprogrammed iPCSs.

**Figure S8: Normalization of read counts in barcode DNA-sequencing datasets for comparing barcode overlap across reprogramming conditions**

A. Representative examples of datasets generated by performing barcode DNA-sequencing of reprogrammed iPSCs separately for iPSCs formed with OKSM alone versus iPSCs formed with OKSM and LSD1 inhibition. Because OKSM and LSD1 inhibition enables more clones to form iPSCs, when performing barcode DNA-sequencing each clone has a lower fraction of clones. This makes using read counts as a proxy for iPSC colony size difficult. To correct for this issue, we applied two different correction methods. First, more simply, we assumed that very large iPSC colonies appearing in both reprogramming conditions should have similar relative colony sizes based on observing our reprogramming cultures and previous analyses in (Emert et al., 2021). We multiplied the reads per million (RPM) for clones reprogrammed in OKSM with LSD1 inhibition by a correction factor, calculated as correction factor = average(RPM in OKSM alone / RPM in OKSM with LSD1 inhibition) for 5-10 clones having the largest number of combined RPM across reprogramming conditions. Second, in future experiments when preparing reprogrammed iPSCs for barcode DNA-sequencing we added in fixed amounts of clones of known barcode as “spike-ins”, enabling us to convert RPM into an approximate number of cells per clone in each reprogramming condition. We plotted the RPM values for each clone in each reprogramming condition before and after applying each correction factor. Clones forming iPSCs in both conditions (near y = x line) are in black, clones forming iPSCs only with OKSM alone (near x-axis) are in orange, and clones forming iPSCs only with OKSM and LSD1 inhibition (near y-axis) are in blue.

B. We evaluated how fold change in number of colonies for iPSCs formed in OKSM with some perturbation (LSD1 inhibition or DOT1L inhibition) versus iPSCs formed in OKSM alone changed when using different minimum cutoffs for classifying a clone as forming an iPSC colony. Clones under this cutoff may represent some mixture of artifacts from PCR amplification and sequencing, small groups of fibroblasts surviving reprogramming but not forming iPSC colonies, and/or small groups of reprogrammed iPSC with either slow growth or delayed reprogramming. We compared how fold change in the number of colonies formed changed for both of our correction factor approaches described in Figure S8A. We show what cutoff we used for our main analyses and plotted the Venn diagrams of barcode overlap across independent biological replicates. The percentages in the Venn diagrams are of all barcodes detected when using a given colony cutoff.

**Figure S9: Constraint analysis by mixing coefficient and Jensen-Shannon distance within and across clones**

A. We asked whether twins sharing a clone barcode were more transcriptionally similar after reprogramming into iPSCs than equal numbers of randomly sampled barcoded cells using the mixing coefficient described in (Goyal et al., 2021) (see Methods). Higher values of the mixing coefficient indicate a higher similarity in expression profiles of the barcoded cells analyzed. Representative UMAPs for different mixing coefficient values are shown. We calculated mixing coefficients within and across clones by comparing cells reprogrammed into iPSCs on different plates in the same reprogramming conditions (shown for OKSM alone in (A) and for OKSM with LSD1 inhibition in (B)) and in different reprogramming conditions (shown in (C)). The boxes in the pairwise correlation plots are colored by the magnitude of the mixing coefficient; white for values near 0 and a gradient to black for values near 1. For each set of pairwise comparisons, we plotted boxplots comparing the mixing coefficients between twins sharing a clone barcode (on matrix diagonal) and non-twins with different clone barcodes (off matrix diagonal). P-values comparing sample medians were calculated using the Wilcoxon rank sum test.

B. We measured the homogeneity of expression states of constituent cells by measuring the Jensen-Shannon distance between the cluster probability distribution associated with each barcode and a random cluster probability distribution generated from 1000 samples of a matched number of cells. We show two examples of clones with similar numbers of cells but different measured Jensen-Shannon distances. Maintaining the organization provided by UMAP, we plotted all barcoded cells in gray and recolored cells corresponding to each clone of interest in dark blue. Additionally, we plotted bar graphs for observed cluster probability distribution (blue) and the average random cluster probability distribution (gray). Finally, we plotted histograms

demonstrating the distribution of randomized Jensen-Shanon distances between random cluster probability distributions generated from 1000 additional random samples and the average random cluster probability distribution from the bar graphs versus the observed Jensen-Shannon distance between the clone barcode of interest and the same average random cluster probability distribution.
C. We plotted the observed Jensen-Shanon distances for each clone barcode in our dataset (in blue) and the mean +/- 95% confidence intervals for the 1000 additional random samplings (in gray) from the histograms in Figure S9B as a function of the number of cells per clone. As the random Jensen-Shanon distance distribution seems to change with the number of cells, we can only make comparisons across clones within a given number of cells per clone.

**Figure S10: Marker genes and barcode distributions for different UMAP clusters in reprogrammed iPSCs in baseline reprogramming versus reprogramming with LSD1 inhibition**
A. To measure what genes were differentially expressed between iPSCs formed in OKSM alone versus OKSM with LSD1 inhibition, we used Seurat's FindMarkers command on matched twins sharing a clone barcode for all clones present in both reprogramming conditions. We plotted the mean +/- 95% confidence interval for the log2(fold change) values for twins reprogrammed with OKSM alone over twins reprogrammed with OKSM and LSD1 inhibition for n = 26 clone barcodes and selected the top 25 upregulated and downregulated genes by mean log2(fold change).
B. Maintaining the organization provided by UMAP in Figure 5A, we plotted all barcoded cells and recolored cells corresponding to expression levels of a subset of genes, with high expression in red and low expression in blue. *OCT4*, *SOX2*, *NANOG*, *PODXL*, and *DNMT3B* were pluripotency markers identified in the literature while *GAPDH* and *UBC* were used as housekeeping genes. We identified markers associated with clusters 0, 1, 2, and 3 (corresponding to reprogrammed iPSCs in our single-cell RNA-sequencing dataset) using Seurat's FindAllMarkers command, and selected a subset of markers associated with pluripotency and/or specific biological processes.
C. To identify clones forming iPSCs only when reprogrammed with OKSM and LSD1 inhibition (i.e., "LSD1i-dependent clones" in light blue) versus clones forming iPSCs in both reprogramming conditions (i.e., "LSD1i-independent clones" in dark blue), we performed barcode DNA-sequencing separately for each programming condition on the leftover iPSC colonies after using a small fraction (<5%) for single-cell RNA-sequencing in Figure 5A. After converting read counts into number of cells using our spike-in controls (see Methods), we plotted each clone on a scatterplot with number of cells in reprogramming plates with OKSM alone on the x-axis and with number of cells in reprogramming plates with OKSM and LSD1 inhibition on the y-axis. We defined LSD1i-dependent clones (in dark blue) as having a >2-fold change increase in number of cells in OKSM and LSD1 inhibition compared with OKSM alone whereas we defined LSD1-independent clones (in dark gray) as having a <2-fold change difference in number of cells in either direction. We performed a similar analysis using our single-cell RNA-sequencing dataset in which each cell was labeled with the clone of origin in the scatterplot on the right. Only a small fraction of reprogrammed iPSCs were used for single-cell RNA-sequencing, representing a significant subsampling. For our final analyses, we defined LSD1i-dependent clones as those that were along the y-axis in both scatterplots and we defined LSD1i-independent clones as those that were along the y = x in the bulk DNA-sequencing but could be either along the y = x or x-axis in the single-cell RNA-sequencing dataset because of the subsampling.
